# Metabolic clogging of mannose triggers genomic instability via dNTP loss in human cancer cells

**DOI:** 10.1101/2022.10.17.512485

**Authors:** Yoichiro Harada, Yu Mizote, Takehiro Suzuki, Mikako Nishida, Toru Hiratsuka, Ayaka Ueda, Yusuke Imagawa, Kento Maeda, Yuki Ohkawa, Junko Murai, Hudson H. Freeze, Eiji Miyoshi, Shigeki Higashiyama, Heiichiro Udono, Naoshi Dohmae, Hideaki Tahara, Naoyuki Taniguchi

## Abstract

Mannose has anti-cancer activity that inhibits cell proliferation and enhances the efficacy of chemotherapy. How mannose exerts its anti-cancer activity, however, remains poorly understood. Here, using genetically engineered human cancer cells that permit the precise control of mannose metabolic flux, we demonstrate that the large influx of mannose exceeding its metabolic capacity induced metabolic remodeling, leading to the generation of slow-cycling cells with limited deoxyribonucleoside triphosphates (dNTPs). This metabolic remodeling impaired dormant origin firing required to rescue stalled forks by cisplatin, thus exacerbating replication stress. Importantly, pharmacological inhibition of *de novo* dNTP biosynthesis was sufficient to retard cell cycle progression, sensitize cells to cisplatin, and inhibit dormant origin firing, suggesting dNTP loss-induced genomic instability as a central mechanism for the anti-cancer activity of mannose.

## Introduction

In mammals, mannose is a monosaccharide that is essential for life and is synthesized *de novo* from glucose through a glycolysis branch (**fig. S1A**). This process requires the action of mannose phosphate isomerase (MPI), the enzyme that catalyzes the interconversion between fructose-6-phosphate (Fruc-6-P) and mannose-6-phosphate (Man-6-P) (Alton et al., 1998). Man-6-P is further converted to GDP-mannose for the biosynthesis of asparagine-linked glycans (N-glycans) in the endoplasmic reticulum (Harada et al., 2013). In the secretory pathway, extensive trimming of N-glycans by mannosidases generates free mannose, which is secreted from cells and contributes to the extracellular pool of mannose (30–130 μM in blood) (Sharma and Freeze, 2011). The extracellular mannose can be taken up by cells and the salvaged mannose is directly converted to Man-6-P, the majority of which is efficiently directed into glycolysis by the action of MPI for unknown reasons (Ichikawa et al., 2014; Sharma and Freeze, 2011).

The large influx of mannose is known to suppress cell proliferation and enhance the efficacy of chemotherapy, particularly in cancer cells that express low levels of MPI (Gonzalez et al., 2018), although the underlying mechanisms remain poorly understood. It has been known for nearly a century that feeding mannose to honeybees, which are believed to express negligible amounts of MPI (Sols et al., 1960), is lethal to these insects (Staudenmayer, 1939). This is because the intracellular levels of Man-6-P exceed the capacity to metabolize it, and the excess Man-6-P inhibits glucose metabolism and decreases the intracellular ATP pool (DeRossi et al., 2006; Sols et al., 1960). This metabolic deficiency is called honeybee syndrome; however, it is unknown whether this syndrome plays a key role in the anti-cancer activity of mannose, and if so, what metabolic checkpoints are targeted by mannose to trigger its anti-cancer activity. Moreover, it is enigmatic how mannose sensitizes poorly proliferating cancer cells to chemotherapy that is designed to target actively proliferating cells. In the present study, we addressed these two fundamental questions by using *MPI*-knockout (MPI-KO) human cancer cells as a model system for honeybee syndrome.

## Results

### Induction of honeybee syndrome suppresses cell proliferation and increases chemosensitivity

To establish MPI-KO human cancer cells using the CRISPR–Cas9 system, we exploited the mannose auxotrophy and sensitivity observed in MPI-KO mouse embryonic fibroblasts (MPI-KO MEFs) (DeRossi et al., 2006). The addition of a physiological concentration of mannose (50 μM, unchallenged) to culture medium supported the proliferation of MPI-KO MEFs (**fig. S1B**). In contrast, mannose starvation or the addition of a supraphysiological concentration of mannose (5 mM, challenged) suppressed the proliferation of MPI-KO MEFs, but not that of wild-type MEFs (**fig. S1C**). On these bases, we knocked out the *MPI* gene in human fibrosarcoma HT1080 cells and screened the gene-edited clones under mannose-unchallenged conditions. Three MPI-KO HT1080 clones were obtained (**Fig. 1A**). In one clone (#1), cell division stopped under mannose-unchallenged conditions, but the other two clones (#2 and #3) could proliferate. These two clones exhibited mannose auxotrophy and sensitivity as expected (**Fig. 1B and fig. S2A**), while the parental HT1080 cells showed marginal defects in cell proliferation at mannose concentrations higher than 15 mM (**Fig. 1C**). The effects of mannose starvation and mannose challenge on the proliferation of MPI-KO cells were almost fully rescued by reintroduction of the human *MPI* gene (**Fig. 1D−F and fig. S2B-D**), ruling out the potential off-target effects of gene editing. As expected, mannose challenge to MPI-KO HT1080 cells caused the dramatic accumulation of hexose-6-phosphate (the sum of glucose-6-phosphate and Man-6-P) compared with that under mannose-unchallenged conditions (**Fig. 1G**). Consistent with the essential role of exogenous mannose in GDP-mannose production for N-glycan biosynthesis in MPI-KO MEFs (Harada et al., 2013), mannose starvation severely decreased N-glycosylation in MPI-KO HT1080 cells (**Fig. 1H**). However, mannose challenge showed negligible effects on N-glycosylation, indicating that mannose challenge suppresses cell proliferation through a mechanism distinct from N-glycosylation defects.

**Fig. 1.**
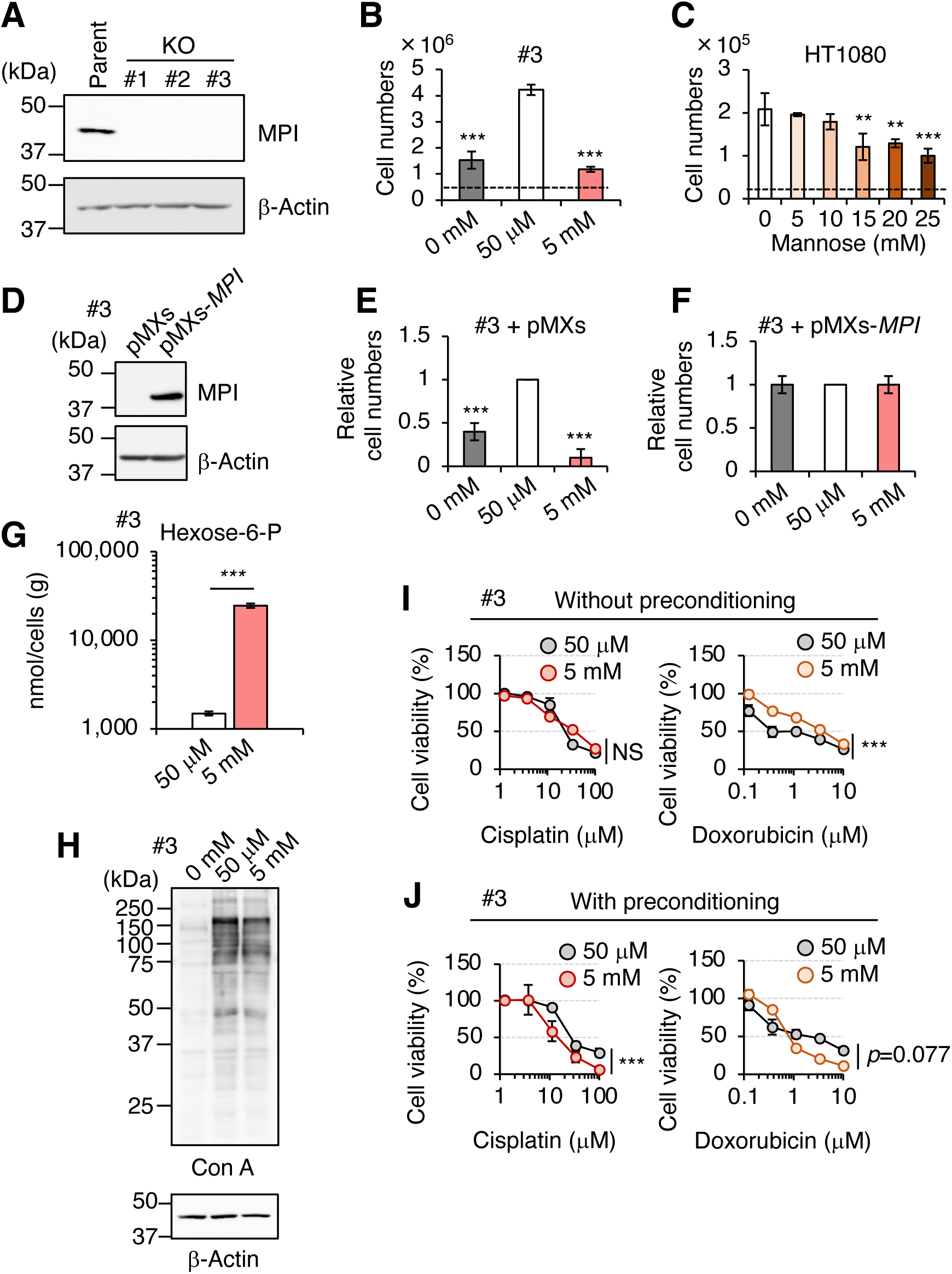
The induction of honeybee syndrome suppresses cell proliferation and increases chemosensitivity. (**A**) Western blot analysis of HT1080 (parent) and MPI-KO HT1080 (KO, clone #1-3) cells. The blots have been vertically flipped for presentation purpose. (**B** and **C**) The cell numbers of MPI-KO HT1080 (#3, **B**) and the parental HT1080 (**C**) cells after 48-h incubation in culture medium supplemented with mannose at the indicated concentrations. A dashed line indicates the cell numbers at seeding. (**D**) Western blot analysis of MPI-KO HT1080 (#3) cells retrovirally transduced with empty vector (pMXs) or human *MPI* gene (pMXs-*MPI*). (**E** and **F**) The relative cell numbers of MPI-KO HT1080 (#3) cells retrovirally transduced with empty vector (pMXs, **E**) or human *MPI* gene (pMXs-*MPI*, **F**) after 48-h incubation under mannose-starved (0 mM), mannose-unchallenged (50 μM), or mannose-challenged (5 mM) conditions. (**G**) The quantification of hexose-6-P in MPI-KO HT1080 (#3) cells cultured in the presence of 50 μM or 5 mM mannose. (**H**) Lectin blot and western blot analyses of MPI-KO HT1080 (#3) cells cultured as in **B**. Con A, concanavalin A lectin. (**I**) Cell viability assay in MPI-KO HT1080 (#3) cells co-treated with mannose (50 μM or 5 mM) and DNA replication inhibitors (cisplatin or doxorubicin) for 24 h (**I**, without preconditioning), or preconditioned with mannose (50 μM or 5 mM) for 24 h, followed by incubation with the DNA replication inhibitors for an additional 24 h (**J**, with preconditioning). Data represent the mean ± SD; n = 3 independent experiments. \**p* < 0.05, \*\**p* < 0.01, and \*\*\**p* < 0.001, NS, not significant, one-way ANOVA with post hoc Dunnett’s test (**B**, **C**, **E**, and **F**), Welch’s t-test (**G**), or two-way ANOVA (**I** and **J**). **Figure 1-source data 1. Original blot images depicting cropped regions for Figure 1A.** **Figure 1-source data 2. Original blot images depicting cropped regions for Figure 1D.** **Figure 1-source data 3. Original blot images depicting cropped regions for Figure 1H.** **Figure 1-figure supplement 1-source data 1. Original blot images depicting cropped regions for Figure S2B.** **Figure 1-figure supplement 2-source data 1. Original blot images depicting cropped regions for Figure S3A.** **Figure 1-figure supplement 2-source data 2. Original blot images depicting cropped regions for Figure S3D.** **Figure 1-figure supplement 2-source data 3. Original blot images depicting cropped regions for Figure S3E.**

Mannose challenge increased the sensitivity of MPI-KO HT1080 cells to DNA replication inhibitors (*i.e.*, cisplatin and doxorubicin) when the cells had been preconditioned with excess mannose prior to the drug treatment (**Fig. 1I, J and fig. S2E, F**). We also generated MPI-KO HeLa cells as another cell model (**fig. S3A-I**), and found that mannose challenge also increased the sensitivity of these cells to cisplatin and doxorubicin (**fig. S3J, K**). All of these results demonstrate that the induction of honeybee syndrome suppresses cell proliferation and increases chemosensitivity in our MPI-KO human cancer cell models. We mostly used MPI-KO HT1080 (#3) cells for subsequent study, while the other cell models showed similar results.

### Mannose challenge generates slow-cycling cells

Cell proliferation is tightly controlled by cell cycle progression. To explore the mechanism behind the anti-proliferative activity of mannose, we compared the cell cycle progression between mannose-challenged and unchallenged MPI-KO HT1080 cells by using a two-color fluorescent ubiquitination-based cell cycle indicator [Fucci(CA)] (Sakaue-Sawano et al., 2017). In this reporter system, G_1_ phase was defined by the exclusive expression of mCherry-hCdt1(1/100)Cy(−), which was rapidly turned off upon the onset of S phase where mVenus-hGem(1/110) gradually accumulated (**Fig. 2A, B and movie S1**). The mCherry-hCdt1(1/100)Cy(−) was re-expressed upon the onset of G_2_ phase (the double-positive phase), followed by the termination of M phase where mVenus-hGem(1/110) was abruptly turned off. Under mannose-unchallenged conditions, MPI-KO HT1080 cells showed exponential growth (**Fig. 2C**) and a typical Fucci(CA) signal profile (**Fig. 2B**), as reported in HeLa cells (Sakaue-Sawano et al., 2017). In contrast, mannose challenge almost completely suppressed cell proliferation (**Fig. 2C**) and significantly prolonged the cell cycle, showing a variety of atypical Fucci(CA) signal profiles (**Fig. 2E, F, fig. S4**). We classified these profiles based on the order of expression of Fucci(CA) reporters [mCherry-hCdt1(1/100)Cy(−) as R, mVenus-hGem(1/110) as G, and the double-negative phase as Dn, **fig. S4**]. The classification analysis revealed that a small proportion of the mannose-challenged cells showed normal-like but strikingly extended Fucci(CA) signal profiles (**Fig. 2E, F and fig. S4**, normal-like, 12% of the total). Notably, these normal-like cell populations frequently failed to undergo cytokinesis in M phase and re-entered G_1_ phase without generating two equivalent daughter cells (**movie S2**). Other fractions included cells that were arrested in G_1_ phase (**Fig. 2E, F and fig. S4**, RRR, 26.7%), that showed little to no double-positive phase (*i.e.*, the G_2_-to-M phase) (**Fig. 2F and fig. S4**, RGR, 15.8% and GRG, 9.9%, GDn, 5.9%, RGG, 5.0%, RGDn, 4.0%; 40.6% of total), or that remained in the double-negative phase (**Fig. 2E, F and fig. S4,** Dn, 8.9%), which is normally seen only at the G_1_/S transition (Sakaue-Sawano et al., 2017). These results suggest that mannose challenge severely impairs the entry of the cells into S phase and its progression to mitotic phase. Strikingly, however, switching of the mannose-challenge medium to the mannose-unchallenged medium after long-term mannose challenge (6 days) resulted in robust cell proliferation (**Fig. 2G**), suggesting that some fraction of the mannose-challenged cells had entered a quiescent state upon mannose challenge. Supporting this assumption, mannose challenge induced early accumulation of the cell cycle inhibitors p21 and p27 (Abukhdeir and Park, 2008) (**Fig. 2H**) and decreased cell populations that were actively incorporating 5-bromo-2’-deoxyuridine (BrdU) into DNA (**Fig. 2I, J**). Small proportions of the mannose-challenged cells exhibited DNA content greater than 4n after 2-day mannose challenge, suggesting that these cell populations underwent endoreplication (Edgar and Orr-Weaver, 2001). Collectively, these results indicate that mannose challenge suppresses cell proliferation through a complex mechanism involving extremely slow cell cycle progression.

**Fig. 2.**
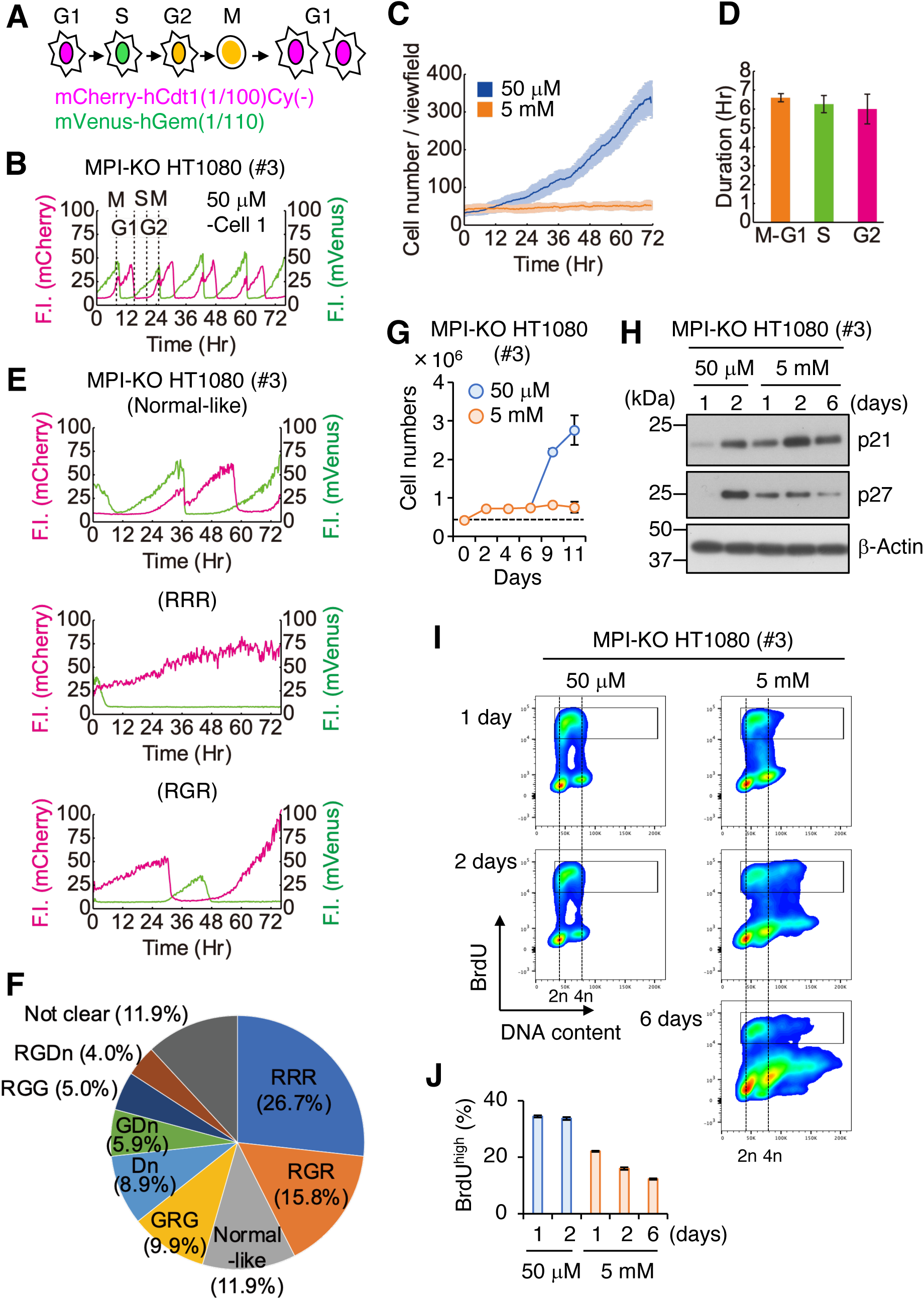
Mannose challenge generates slow-cycling cells. (**A**) Schematic representation of cell cycle progression (G1, S, G2, and M phases) visualized by Fucci(CA). Magenta, green, and yellow indicate the expression of mCherry-hCdt1(1/100)Cy(−), mVenus-hGem(1/110), and both, respectively. (**B**) A representative Fucci(CA) signal profile in MPI-KO HT1080 cells under the mannose-unchallenged conditions. Fluorescent intensity (F.I.) for mCherry (left y-axis) and mVenus (right y-axis) is shown by magenta and green, respectively. Individual phases of the cell cycle (G_1_, S, G_2_, and M) are demarcated by dashed lines. (**C**) The number of MPI-KO HT1080 cells during the time-lapse imaging under mannose-unchallenged (blue) and mannose-challenged (orange) culture conditions. (**D**) The durations of M–G_1_ (the sum of both phases), S, and G_2_ phases (n = 1109 cells) in MPI-KO HT1080 cells under mannose-unchallenged conditions. (**E**) Representative Fucci(CA) signal profiles of MPI-KO HT1080 cells cultured under mannose-challenged conditions for 76 h. The order of expression of mCherry-hCdt1(1/100)Cy(−) (denoted by R) and mVenus-hGem(1/110) (denoted by G) was used to classify the Fucci profiles and the classification is indicated in parentheses. See also fig. S4. (**F**) The proportion of Fucci(CA) signal profiles of the mannose-challenged MPI-KO HT1080 cells. R, mCherry-hCdt1(1/100)Cy(-−) positive; G, mVenus-hGem(1/100) positive; Dn, double negative for Fucci indicators. One hundred cells were visually inspected for the classification and the cells that could not be classified were categorized as “not clear.” See also fig. S4. (**G**) The number of MPI-KO HT1080 (#3) cells. The cells were cultured in the presence of 5 mM mannose for 6 days, and they were further cultured in the presence of 50 μM or 5 mM mannose for 5 days. A dashed line indicates the cell numbers at seeding. (**H**) Western blot analysis of MPI-KO HT1080 (#3) cells cultured in the presence of 50 μM or 5 mM mannose for the indicated time. (**I**) Flow cytometry for BrdU and DNA content (Hoechst33342) in MPI-KO HT1080 (#3) cells cultured in the presence of 50 μM or 5 mM mannose for the indicated time. The cells were labeled with 10 μM BrdU for 1 h before harvest. (**J**) Quantification of BrdU^high^ cells in **I** (in the gated populations). Data represent the mean ± SD; n = 4 independent fields (**C**) and n = 3 independent experiments (**G** and **J**). **Figure 2-source data 1. Original blot images depicting cropped regions for Figure 2H.**

### Mannose challenge limits DNA synthesis at ongoing replication forks

We adopted a proteomic approach to dissect the molecular mechanism by which mannose challenge generated slow-cycling cells in MPI-KO HT1080. Of over 7,000 proteins identified in the proteomic datasets (**data S1**), proteins that were significantly up- and downregulated relative to those in the cells cultured for 1 day under mannose-unchallenged conditions were extracted at each time point (**data S2**). Functional annotation analysis of the proteomic data using DAVID bioinformatic resources (https://david.ncifcrf.gov) revealed the downregulation of proteins related to the cell cycle and DNA replication in mannose-challenged cells (**Fig. 3A**), which were enriched with the mini-chromosome maintenance 2-7 (MCM2-7) complex (**Fig. 3B**). Western blot analysis and quantitative polymerase chain reaction (qPCR) confirmed the decrease in the expression levels of MCM2-7 proteins (**Fig. 3C**) and their genes (**Fig. 3D**) during mannose challenge over 6 days. The MCM2-7 complex is a core component of DNA helicase that unwinds the DNA duplex at replication forks in S phase (Jones et al., 2021; Rzechorzek et al., 2020), and the complex also plays a central role in the licensing of replication origins in G_1_ phase by forming a pre-replicative complex with the six-subunit origin recognition complex (ORC1–6), CDC6, and CDT1 on chromatin (Frigola et al., 2017; Remus et al., 2009; Zhai et al., 2017). CDC6 and CDT1, which were not detected in our proteomic analysis, were depleted after 2-day mannose challenge, whereas the ORC2 subunit was expressed at relatively constant levels (**Fig. 3C**). These results indicate that mannose challenge induces slow proteomic alterations in origin licensing factors, while this cannot be an essential trigger for the generation of slow-cycling cells, as these cells appeared immediately after mannose challenge.

**Fig. 3.**
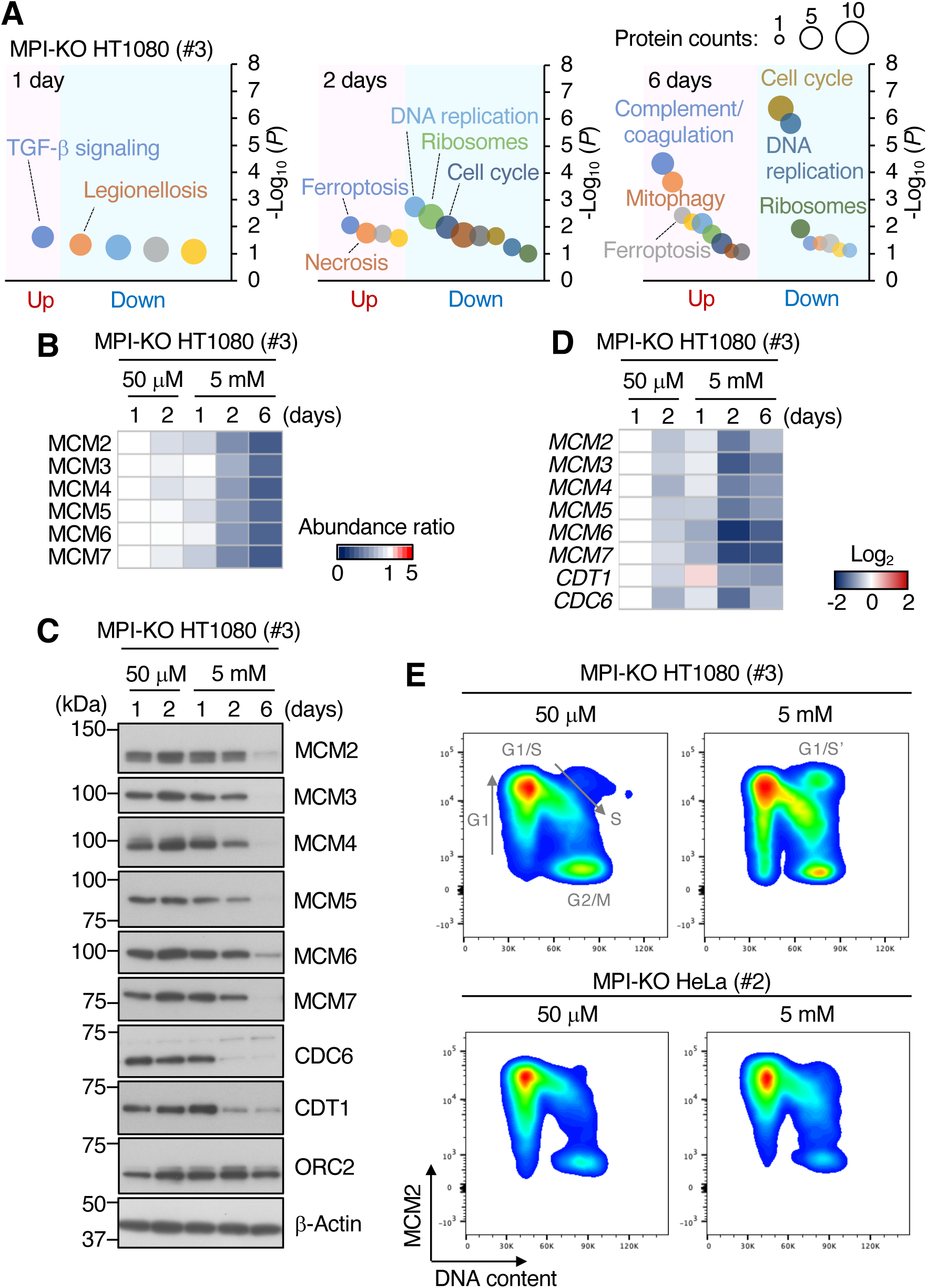
Mannose challenge limits DNA synthesis at ongoing replication forks. (**A**) Functional annotation analysis of the proteomic data in MPI-KO HT1080 (#3) cells cultured in the presence of 50 μM or 5 mM mannose for the indicated time. (**B**) Heatmap representation of the relative amounts of MCM2-7 proteins in MPI-KO HT1080 (#3) cells cultured in the presence of 50 μM or 5 mM mannose for the indicated time. (**C**) Western blot analysis of whole-cell lysates of MPI-KO HT1080 (#3) cells cultured in the presence of 50 μM or 5 mM mannose for the indicated time. (**D**) Heatmap representation of the relative expression levels of *MCM2*, *MCM3*, *MCM4*, *MCM5*, *MCM6*, *MCM7*, *CDT1*, and *CDC6* genes in MPI-KO HT1080 (#3) cells cultured in the presence of 50 μM or 5 mM mannose for the indicated time. (**E**) Chromatin flow cytometry for MCM2 and DNA content (Hoechst33342) in MPI-KO HT1080 (#3) or MPI-KO HeLa (#2) cells cultured in the presence of 50 μM or 5 mM mannose for 24 h. Progression of cell cycle (G_1_, G_1_/S boundary, S, and G_2_/M phases) is indicated by arrows. G_1_/S’ denotes the G_1_/S boundary of cell populations that underwent endoreplication. **Figure 3-source data 1. Original blot images depicting cropped regions for Figure 3C.** **Figure 3-source data 2. Original blot images depicting cropped regions for Figure 3C.** **Figure 3-source data 3. Original blot images depicting cropped regions for Figure 3C.** **Figure 3-source data 4. Original blot images depicting cropped regions for Figure 3C.**

We performed chromatin flow cytometry for MCM2 to elucidate the impact of mannose challenge on the chromatin-bound states of the MCM complex. Under the mannose-unchallenged conditions, the binding of MCM2 to chromatin peaked at the G_1_/S boundary in MPI-KO HT1080 cells and MPI-KO HeLa cells (**Fig. 3E**), indicating that replication origins were fully licensed (Matson et al., 2017). The cells with the fully licensed origins entered S phase as a tight population, and MCM2 dissociated from chromatin as two replication forks converged and terminated (**Fig. 3E**) (Low et al., 2020). Mannose challenge did not severely impair origin licensing (**Fig. 3E**), but the same treatment caused the accumulation of MCM2-positive chromatin in S phase (**Fig. 3E**). Notably, the mannose-challenged cells were not actively incorporating BrdU into DNA (**Fig. 2I, J and fig. S5A**), suggesting that ongoing replication forks were stuck on chromatin with little DNA synthesis under the mannose-challenged conditions. Taking these findings together, the limited DNA synthesis at ongoing replication forks likely contributes to the abnormally extended progression of S phase in the mannose-challenged cells.

### Mannose challenge disengages dormant origins from DNA synthesis during replication stress

Although our findings indicated that mannose challenge limits DNA synthesis at ongoing replication forks, its relevance to the increased chemosensitivity was still unclear. In humans, replication origins are licensed far more than actually used for DNA replication (Langley et al., 2016). The excess origins remain dormant under physiological conditions, while the dormant origins are activated for DNA replication when nearby replication forks stall upon encountering DNA lesions, thus preventing cells from the permanent replication arrest that leads to cell death (Blow et al., 2011; Ge et al., 2007; Kawabata et al., 2011; Shima et al., 2007). To test whether mannose challenge may also impair DNA synthesis from dormant origins during replication stress, we compared BrdU incorporation between the cells that were first pulsed with cisplatin to induce replication stress, followed by being left untreated or being treated with an inhibitor of the ataxia telangiectasia and Rad3-related protein (ATR) to forcibly activate dormant origins (Moiseeva and Bakkenist, 2019; Moiseeva et al., 2019) (**Fig. 4A**). In the mannose-unchallenged MPI-KO HT1080 cells (**Fig. 4B, C**) and MPI-KO HeLa cells (**fig. S5A, B**), the cisplatin treatment alone partially suppressed the incorporation of BrdU, which was greatly recovered by ATR inhibition with VE-821 (ATRi) (Charrier et al., 2011; Prevo et al., 2012; Reaper et al., 2011), indicating that dormant origins are present in excess and their activation can engage in DNA synthesis during replication stress. In the mannose-challenged cells, however, the cisplatin treatment more severely reduced BrdU incorporation, which was barely restored by ATRi treatment (**Fig. 4B, C and fig. S5A, B**). Forced activation of dormant origins during replication stress is known to cause the uncoupling of DNA unwinding and synthesis (Murai et al., 2018), leading to the accumulation of single-stranded DNA on chromatin marked by phosphorylation of the replication protein A2 (RPA2) (**Fig. 4D**), confirming the activation of dormant origins by ATRi treatment in both mannose-challenged and -unchallenged cells. These results indicate that mannose challenge limits DNA synthesis from dormant origins during replication stress.

**Fig. 4.**
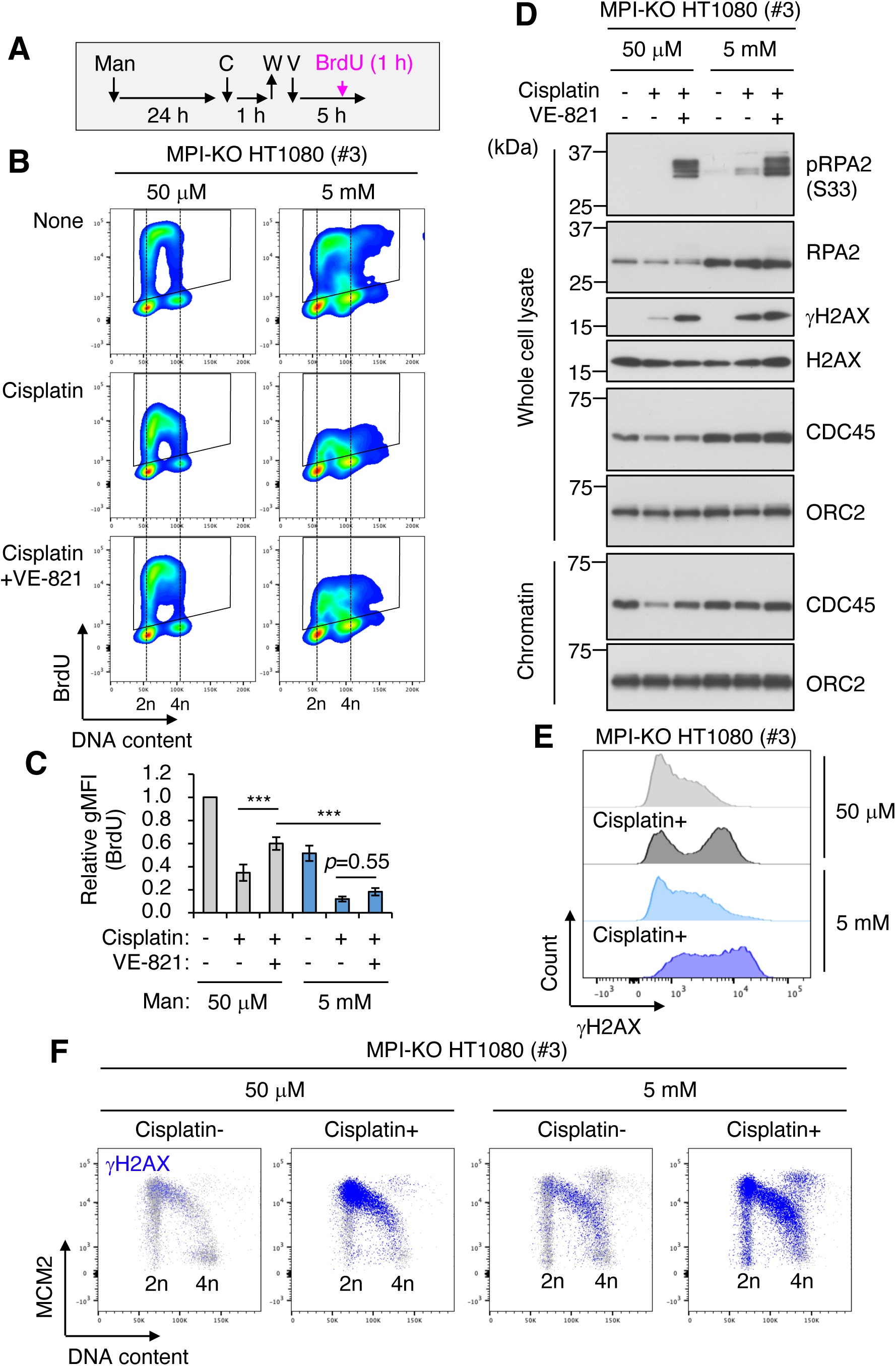
Mannose challenge disengages dormant origins from DNA synthesis during replication stress. (**A**) Schematic representation of drug treatments. Man, 50 μM or 5 mM mannose; C, 100 μM cisplatin; W, wash; V, 1 μM VE-821; BrdU, 10 μM 5-bromo-2′-deoxyuridine. (**B**) Flow cytometry for BrdU and DNA content (Hoechst33342) in MPI-KO HT1080 (#3) cells treated as indicated in **A**. (**C**) Relative geometric mean fluorescent intensity (gMFI) of BrdU in **B** (in the gated populations). (**D**) Western blot analysis of whole-cell lysates and chromatin fractions of MPI-KO HT1080 (#3) cells treated as indicated in **A**, except that BrdU labeling was omitted. (**E** and **F**) Chromatin flow cytometry for γH2AX, MCM2, and DNA content (Hoechst33342) in MPI-KO HT1080 (#3) cells treated as in **A**, except that VE-821 treatment and BrdU labeling were omitted. Data represent the mean ± SD; n = 3 independent experiments. \*\*\**p* < 0.001, one-way ANOVA with post hoc Tukey’s test (**C**). **Figure 4-source data 1. Original blot images depicting cropped regions for Figure 4D.** **Figure 4-source data 2. Original blot images depicting cropped regions for Figure 4D.** **Figure 4-source data 3. Original blot images depicting cropped regions for Figure 4D.** **Figure 4-source data 4. Original blot images depicting cropped regions for Figure 4D.**

The replication initiation step requires chromatin loading of cell division cycle 45 (CDC45), one of the essential and limiting factors for replication initiation (Moyer et al., 2006), while this process was not severely impaired by mannose challenge (**Fig. 4D and fig. S6A, B**). Under mannose-unchallenged conditions, cisplatin treatment induced the unloading of CDC45 from chromatin, which was restored by the forced activation of dormant origins (**Fig. 4D and fig. S6A, B**). However, mannose challenge abrogated the CDC45 unloading/reloading dynamics under replication stress conditions (**Fig. 4D and fig. S6A, B**). The deficiency in the DNA synthesis from dormant origins and the CDC45 dynamics was associated with the increase in cisplatin-induced γH2AX (**Fig. 4D, E and fig. S5C**), the phosphorylated form of the histone H2AX that marks DNA double-strand breaks (Kuo and Yang, 2008). The endogenous levels of γH2AX were detected in S phase of both mannose-challenged and -unchallenged cells (**Fig. 4F, fig. S5D and S6C**), and cisplatin strongly induced γH2AX throughout S phase even in the mannose-challenged cells (**Fig. 4F and fig. S5D**). Together, these results indicate that mannose challenge disengages dormant origins from DNA synthesis during replication stress, thus exacerbating DNA damage.

### Mannose challenge generates a distinct metabolic landscape

Although our data showed that mannose challenge generated slow-cycling cells that failed to engage dormant origins in DNA synthesis during replication stress, the underlying mechanism that functionally links these two phenotypes was still unclear. Since cell cycle progression and DNA replication are metabolically demanding processes (Heinemann and Zylstra, 2022), we hypothesized that the metabolic checkpoints activated by honeybee syndrome (DeRossi et al., 2006; Sols et al., 1960) may play a key role in the anti-cancer activity of mannose. To test this hypothesis, we first examined the impact of mannose challenge on the cellular bioenergetics in MPI-KO HT1080 cells by monitoring the real-time changes in glycolytic flux in the form of extracellular acidification rate (ECAR) and the activity of oxidative phosphorylation (OXPHOS) in the form of oxygen consumption rate (OCR). Mannose challenge increased OCR in response to a steep drop of ECAR (**Fig. 5A, B**). This bioenergetic compensation was not sufficient to maintain the ATP production rates in full (**Fig. 5C**). Consistent with this, mannose challenge decreased the ATP pool in MPI-KO HT1080 cells, MPI-KO HeLa cells, and MPI-KO MEFs (**Fig. 5D, E and fig. S7**), and the remaining pool was completely depleted by co-treatment with IACS-010759, a preclinical small-molecule inhibitor of complex I of the mitochondrial respiratory chain (Molina et al., 2018) (**Fig. 5D, E and fig. S7**), which in turn increased necrosis (**Fig. 5F**). These results indicate that, in honeybee syndrome, the bioenergetic balance is impaired by glycolysis being acutely shut down and OXPHOS being boosted.

**Fig. 5.**
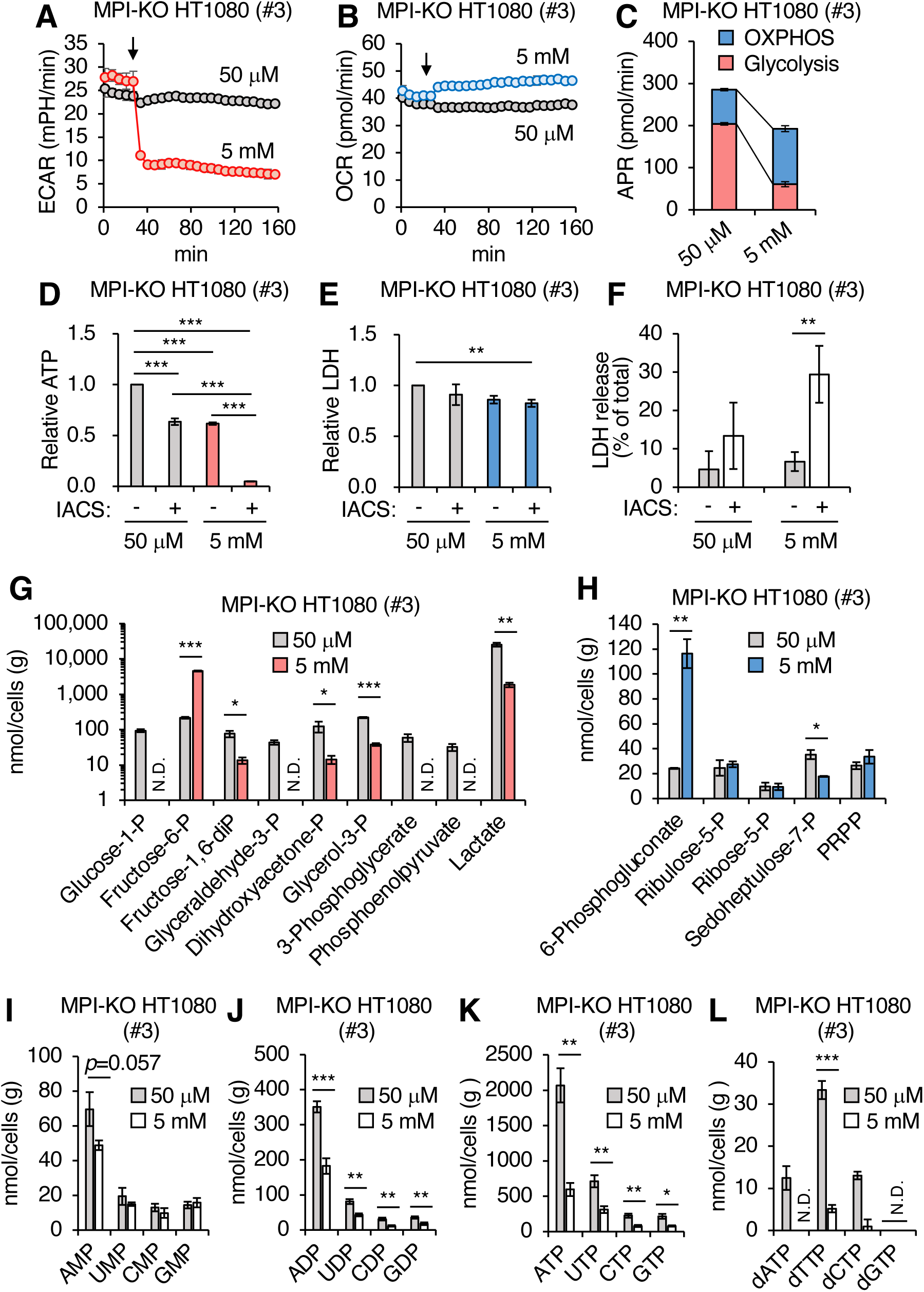
Mannose challenge generates a distinct metabolic landscape. (**A** and **B**) Extracellular acidification rates (ECAR, **A**) and oxygen consumption rates (OCR, **B**) in MPI-KO HT1080 (#3) cells. The cells were first cultured in 50 μM mannose, and then the medium alone or mannose (5 mM final) was added to the cells at the timing indicated by the downward arrows. (**C**) ATP production rates (APR) estimated from the data in **A** and **B**. See also the Materials and Methods. (**D**) Relative ATP levels in MPI-KO HT1080 (#3) cells treated for 6 h with or without 30 nM IACS-010759 (IACS) in the presence of 50 μM or 5 mM mannose. (**E**) Relative lactate dehydrogenase (LDH) levels in MPI-KO HT1080 (#3) cells treated as in **D**. (**F**) LDH release from MPI-KO HT1080 (#3) cells treated for 24 h with or without 30 nM IACS-010759 (IACS) in the presence of 50 μM or 5 mM mannose. (**G**–**L**) Metabolomic profiling of glycolysis (**G**), pentose phosphate pathway (**H**), nucleotides (**I**–**K**), and deoxynucleoside triphosphates (**L**) in MPI-KO HT1080 (#3) cells cultured for 24 h in the presence of 50 μM or 5 mM mannose. P, phosphate; PRPP, 5-phosphoribosyl-1-pyrophosphate. Data represent the mean ± SD; n = 3 independent experiments. \**p* < 0.05, \*\**p* < 0.01, and \*\*\**p* < 0.001, one-way ANOVA with post hoc Tukey’s test (**D**–**F**) or Welch’s t-test (**G**–**L**).

To provide deeper insights into the metabolic landscape generated by honeybee syndrome, we compared the metabolome between mannose-challenged and -unchallenged MPI-KO HT1080 cells (**fig. S8A** and **data S3**). One-day mannose challenge greatly increased Fruc-6-P, while the three-carbon metabolites in the lower glycolysis chain, including lactate, were substantially decreased (**Fig. 5G**), supporting our earlier findings that mannose challenge decreased ECAR to the basal level. Despite this glycolytic insufficiency, the amounts of metabolites in oxidative and non-oxidative arms of the pentose phosphate pathway (PPP) remained relatively unchanged in the mannose-challenged cells, except for the large accumulation of 6-phosphogluconoate and the slight decrease in sedoheptulose-7-phosphate (**Fig. 5H**). In contrast, mannose challenge severely decreased tricarboxylic acid (TCA) cycle intermediates (isocitrate, 2-oxoglutarate, succinate, fumarate, and malate) that are used to generate NADH and FADH_2_ as electron donors for the mitochondrial respiratory chain (**fig. S8B**), suggesting their large consumption by the boosting of OXPHOS in honeybee syndrome. Strikingly, the amounts of nucleoside diphosphates and triphosphates, but not nucleoside monophosphates, were proportionally and moderately decreased (**Fig. 5I−K**), whereas deoxyribonucleoside triphosphates (dNTPs) were substantially decreased in the mannose-challenged cells (**Fig. 5L**). Collectively, these results indicate that mannose challenge generates a distinct metabolic landscape in MPI-KO HT1080 cells, which ultimately lead to the depletion of dNTP pools.

### Pharmacological inhibition of *de novo* dNTP biosynthesis retards cell cycle progression, increases chemosensitivity, and inhibits DNA synthesis from dormant origins

dNTPs are the essential donor substrates for DNA synthesis, raising the possibility that the dNTP loss caused by mannose challenge may be a major mechanism linking the generation of slow-cycling cells and the failure to engage dormant origins in DNA synthesis during replication stress. To test this hypothesis, we inhibited *de novo* synthesis of dNTPs with hydroxyurea (HU), a highly potent inhibitor of ribonucleotide reductase (RNR) (Elford, 1968), and found that the combination of HU and cisplatin more severely reduced cell viability than cisplatin treatment alone, independently of the presence or absence of the *MPI* gene (**Fig. 6A**). This chemosensitizing effect of HU was associated with the strong induction of γH2AX (**Fig. 6B**). HU is widely used as an agent to arrest cells in S phase by depleting dNTP (Davis et al., 2001). Profiling of the chromatin-bound MCM2 showed that HU treatment at a moderate concentration (0.25 mM) resulted in the arrest of cells in the early to late S phase, while a higher concentration of HU (1 mM) almost completely arrested the cells at the G_1_/S boundary (**Fig. 6C**), partly recapitulating the inhibitory effects of mannose challenge on cell cycle progression. As with the case of mannose challenge, the forced activation of dormant origins by ATRi failed to increase BrdU uptake in the presence of HU (**Fig. 6D–G**), indicating that the DNA synthesis from dormant origins is highly sensitive to the pool size of dNTPs. Unlike mannose challenge, however, HU treatment did not severely impair the CDC45 unloading/reloading dynamics during replication stress (**Fig. 6G**), indicating that the insufficient dNTP pool is a major cause of the disengagement of dormant origins from DNA synthesis.

**Fig. 6.**
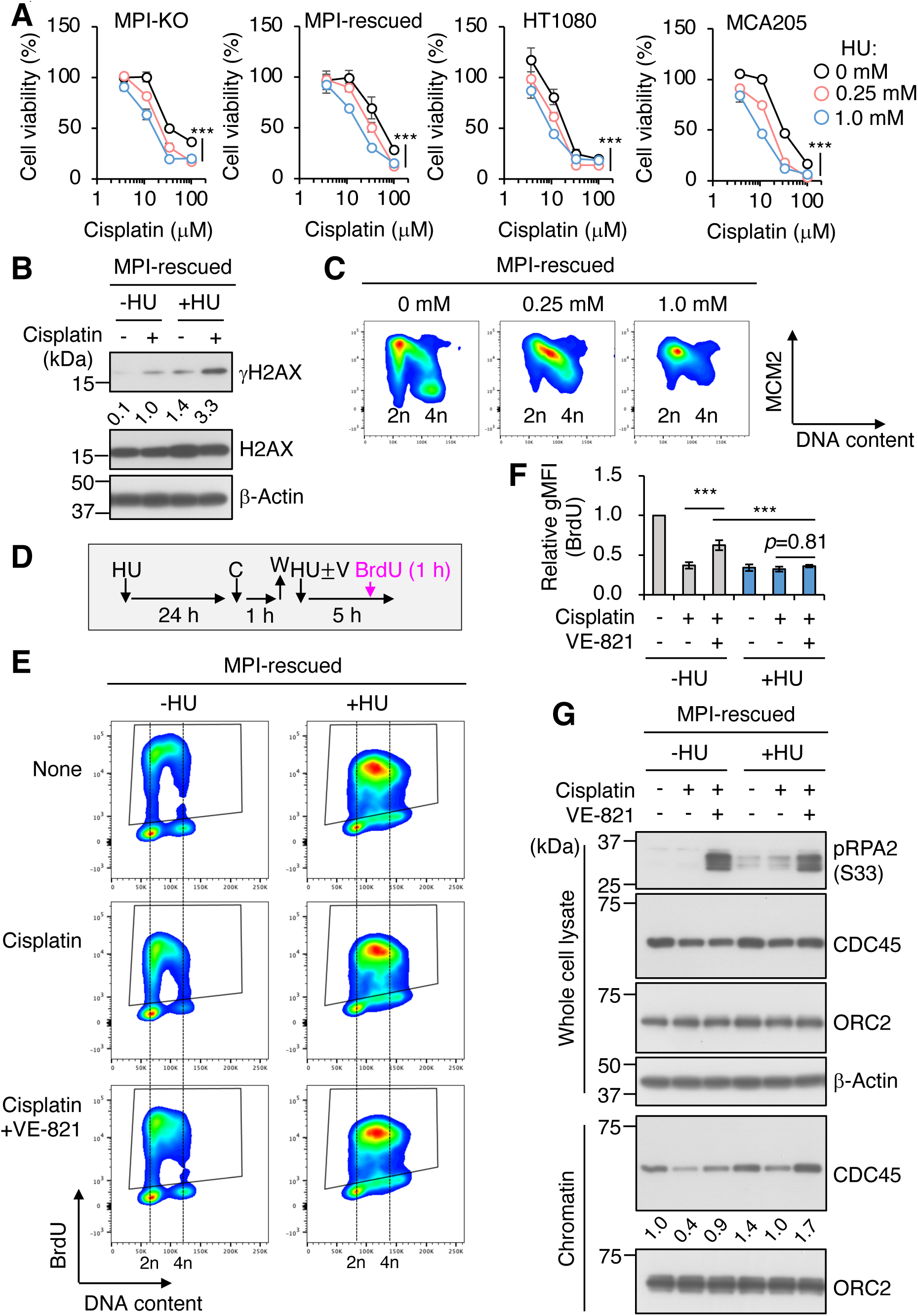
Pharmacological inhibition of *de novo* dNTP biosynthesis retards cell cycle progression, increases chemosensitivity, and inhibits DNA synthesis from dormant origins. (**A**) Cell viability assay in MPI-KO HT1080 (#3), the *MPI*-rescued cells (pMXs-*MPI*), the parental HT1080, and mouse fibrosarcoma MCA205 cells that were preconditioned with hydroxyurea (HU) for 24 h, followed by incubation with cisplatin for an additional 24 h at the indicated concentrations. (**B**) Western blot analysis of whole-cell lysates from the *MPI*-rescued MPI-KO HT1080 (#3) cells preconditioned with 0.25 mM HU for 24 h and pulsed with 100 μM cisplatin for 1 h, followed by a 5-h chase. Numbers indicate the relative amounts of γH2AX (the average of two independent experiments each from the *MPI*-rescued MPI-KO HT1080 cells and the parental HT1080 cells). (**C**) Chromatin flow cytometry for MCM2 and DNA content (Hoechst33342) in the *MPI*-rescued MPI-KO HT1080 (#3) cells treated with or without HU for 24 h at the indicated concentrations. (**D**) Schematic representation of drug treatment. HU, 0.25 mM hydroxyurea; C, 100 μM cisplatin; W, wash; V, 1 μM VE-821; 10 μM BrdU, 5-bromo-2′-deoxyuridine. (**E**) Flow cytometry for BrdU and DNA content (Hoechst33342) in the *MPI*-rescued MPI-KO HT1080 (#3) cells that were treated as indicated in D. (**F**) Relative geometric mean fluorescent intensity (gMFI) of BrdU in D (in the gated populations). (**G**) Western blot analysis of whole-cell lysates and chromatin fractions of the *MPI*-rescued MPI-KO HT1080 (#3) cells treated as indicated in D, except that BrdU labeling was omitted. Numbers indicate the relative amounts of chromatin-bound CDC45 (the average of three independent experiments). Data represent the mean ± SD; n = 3 independent experiments. \**p* < 0.05, \*\**p* < 0.01, and \*\*\**p* < 0.001, two-way ANOVA (A) or one-way ANOVA (E) with post hoc Tukey’s test. **Figure 6-source data 1. Original blot images depicting cropped regions for Figure 6B.** **Figure 6-source data 2. Original blot images depicting cropped regions for Figure 6G.** **Figure 6-source data 3. Original blot images depicting cropped regions for Figure 6G.** **Figure 6-source data 4. Original blot images depicting cropped regions for Figure 6G.**

## Discussion

MPI is the sole enzyme that catalyzes the interconversion between Fruc-6-P and Man-6-P in mammals. The conversion of Fruc-6-P to Man-6-P mediated by MPI is central to the synthesis of GDP-mannose from abundant glucose for normal glycosylation. In contrast, the conversion of Man-6-P to Fruc-6-P mediated by the same enzyme is critical to directing the excess Man-6-P to glycolysis, while the cellular function of this seemingly wasteful metabolic pathway has long remained unknown. In the present study, we employed MPI-KO human cancer cells to explore the key mechanism behind the anti-cancer activity of mannose, and demonstrated that the large influx of mannose exceeding the capacity to metabolize it, that is, the onset of honeybee syndrome, induced dramatic metabolic remodeling that led to dNTP loss. These cells were cycling extremely slowly, and upon encountering replication stress, they were unable to rescue stalled forks via dormant origins, thus exacerbating replication stress. These findings shed light on the conversion of Man-6-P to Fruc-6-P mediated by MPI as a genome guard through the maintenance of dNTP homeostasis, which could be threatened by dietary mannose intake.

Our findings demonstrate that the metabolic landscape formed by honeybee syndrome is far more complex than previously proposed (DeRossi et al., 2006; Sols et al., 1960). Acute shutdown of glycolysis is the hallmark of honeybee syndrome, which concomitantly boosts OXPHOS and forces the cellular metabolome to adapt to this bioenergetic imbalance. Strikingly, the metabolic checkpoints targeted in honeybee syndrome are quite distinct from those expected from *in vitro* enzymatic activity assays (DeRossi et al., 2006). In particular, despite the fact that Man-6-P is a highly potent inhibitor of glucose-6-phosphate dehydrogenase *in vitro* (DeRossi et al., 2006), PPP metabolites were shown not to be depleted in honeybee syndrome, suggesting that previously unidentified mechanisms are operating to cause dNTP loss. In addition to the essential role of dNTPs in the normal progression of S phase, accumulating evidence that the temporal regulation of glycolysis and OXPHOS drives the G_1_/S transition and chromosome segregation (Icard et al., 2019; Salazar-Roa and Malumbres, 2017) supports our hypothesis that the unscheduled biogenetic shift, together with dNTP loss caused by honeybee syndrome, globally impairs cell cycle progression. Taking the findings together, the massive changes in crosstalk between metabolic networks and cell cycle machinery in honeybee syndrome potentially contribute to the generation of slow-cycling cells.

Given the homeostatic role and ubiquitous expression of MPI (Proudfoot et al., 1994), supraphysiological concentrations of mannose are required to trigger its anti-cancer activity in a wide variety of cancer cells (Gonzalez et al., 2018; Zhang et al., 2022). This limitation may hamper the possible clinical use of mannose in cancer therapy. Here, we provide experimental evidence showing that pharmacological inhibition of *de novo* dNTP synthesis, which is critical for the recovery from replication stress (Hakansson et al., 2006; Tanaka et al., 2000), partly mimics the anti-cancer activity of mannose and chemosensitizes cells to cisplatin independently of the MPI expression and mannose dosing. Our finding that DNA synthesis from dormant origins during replication stress is highly sensitive to the dNTP pool size is in good agreement with the therapeutic advantages of RNR inhibition in enhancing the efficacy of radiochemotherapy (Kunos and Ivy, 2018).

In conclusion, we demonstrate that honeybee syndrome plays a critical role in the anti-cancer activity of mannose, at least in part through dNTP loss. Ensuring appropriate amounts of dNTP is essential for normal DNA replication, as well as for the recovery from replication stress, thus providing a plausible mechanism for how mannose sensitizes poorly proliferating cancer cells to chemotherapy.

## Materials and Methods

### Experimental models

The human fibrosarcoma cell line HT1080 used in this study was our laboratory stock and validated by short tandem repeat profiling (Promega). The human cervix adenocarcinoma cell line HeLa was obtained from RIKEN BioResource Research Center. The mouse fibrosarcoma cell line MCA205 was a kind gift from Dr. S. A. Rosenberg (National Cancer Institute, Bethesda, MD). These cell lines were cultured in Dulbecco’s modified Eagle’s medium (DMEM; 045-30285, FUJIFILM Wako) supplemented with 4 mM glutamine (073-05391, FUJIFILM Wako) and 10% fetal bovine serum (FBS) (complete DMEM) at 37°C in a 5% CO_2_ atmosphere.

Immortalized wild-type MEFs and MPI-KO MEFs were prepared in a previous study (DeRossi et al., 2006). MPI-KO HT1080 cells and MPI-KO HeLa cells were established in this study (see below). All MPI-KO cell lines were maintained in complete DMEM supplemented with mannose (Man; 130-00872, FUJIFILM Wako) at a concentration of 20 μM at 37°C in a 5% (for MPI-KO HT1080 cells and MPI-KO HeLa cells) and 10% (for MEFs) CO_2_ atmosphere. Trypan blue dye (0.4%, w/v, FUJIFILM Wako) was mixed with equal volumes of cell suspension for cell counting.

### Cell treatments

Cells were treated with 0−100 μM cisplatin (D3371, Tokyo Chemical Industry), 0−10 μM doxorubicin (040-21521, FUJIFILM Wako), 30 nM IACS-010759 (S8731, Selleckchem), 1 μM VE-821 (SML1415, Sigma Aldrich), or 0−1 mM HU (085-06653, FUJIFILM Wako) for the indicated time. As a negative control, cells were treated with vehicle alone.

### Gene editing

*MPI* gene was knocked out by CRISPR–Cas9 gene editing using the Edit-R system (Dharmacon), in accordance with the manufacturer’s instructions. Briefly, HT1080 cells or HeLa cells were seeded and cultured for 24 h prior to transfection. Synthetic CRISPR RNAs (CM-011729-01 and CM-011729-03, Dharmacon; 10 μM, 2 μL each) targeting the *MPI* gene were mixed with 4 μL of 10 μM trans-activating CRISPR RNA (tracrRNA; U-002005-05, Dharmacon) and transfected with 1 μg of Edit-R SMART Cas9_mCMV_(PuroR) expression plasmid (U-005200-120, Dharmacon) using 5 μL of Lipofectamine 2000 (11668027, Thermo Fisher Scientific) in 400 μL of Opti-MEM (31985062, Thermo Fisher Scientific). Transfectants were selected in complete medium supplemented with 50 μM Man and 1 μg/mL puromycin, and then cloned by limiting dilution. For HT1080 cells, MPI-KO clones were screened by polymerase chain reaction (PCR) using genomic DNA as a template and two primer sets (Supplementary Table S1) for amplifying the wild-type allele and for the KO allele of the *MPI* locus. The PCR products for the KO allele were subjected to Sanger sequencing. For HeLa cells, MPI-KO clones were screened by mannose auxotrophy and sensitivity, as well as Western blotting using anti-MPI antibody (see below).

### Preparation of cDNA

Total RNA was prepared using TRIzol Reagent (15596026, Thermo Fisher Scientific) or RNeasy Mini Kit (74106, Qiagen), in accordance with the manufacturer’s instructions. cDNA was prepared by reverse transcription using total RNA (4 μg), oligo dT primers, and SuperScript IV Reverse Transcriptase (18090010, Thermo Fisher Scientific), in accordance with the manufacturer’s instructions.

### Preparation of plasmids

The plasmids used in this study are listed in Supplementary Table S2. The coding sequence of the human *MPI* gene was amplified by PCR using Phusion High-Fidelity DNA polymerase (M0530, New England BioLabs), a primer set (Supplementary Table S1), and cDNA from HuH-7 cells as a template. The PCR products were purified from agarose gels using NucleoSpin Gel and PCR Clean-Up Kit (740609, Clontech) and subcloned into the pENTR/D-TOPO vector (K240020, Thermo Fisher Scientific), in accordance with the manufacturer’s instructions. The yielded plasmid (pENTR-hMPI) was used as a template to amplify the *MPI* gene by PCR using a primer set (Supplementary Table S1) and the PCR products were cloned into pMXs-Neo retroviral expression vector (RTV-011, Cell Biolabs) to yield pMXs-Neo-hMPI by using In-Fusion HD Cloning Kit (639648, Clontech), in accordance with the manufacturer’s instructions. mCherry-hCdt1(1/100)Cy(−)/pcDNA3 was purchased from RIKEN BioResource Research Center. The coding sequence of mCherry-hCdt1(1/100)Cy(−) was amplified by PCR using a primer set (Supplementary Table S1) and cloned into pMXs-Neo to yield pMXs-Neo-mCherry-hCdt1(1/100)Cy(−) using the In-Fusion HD Cloning Kit. pMMLV-mVenus-hGem(1/110):IRES:Bsd was constructed and purchased from Vector Builder.

### Retroviral packaging and transduction

The Plat-A retroviral packaging cell line (1 × 10^6^ cells, Cell Biolabs) was seeded on collagen I-coated six-well plates (4810-010, IWAKI) and cultured in complete medium for 24 h. After the culture medium had been replaced with 1 mL of fresh complete medium, 1 mL of Opti-MEM containing 3 μg of plasmids (Supplementary Table S2), 9 μL of Lipofectamine 3000, and 6 μL of P3000 reagent (L3000015, Thermo Fisher Scientific) was added to the cells, followed by culture for 48 h. The culture medium was then passed through a 0.80 μm syringe filter (SLAA033SS, Merck Millipore). The filtrate, which contained retroviral particles, was mixed with hexadimethrine bromide (8 μg/mL; 17736-44, Nacalai Tesque) and Man (20 μM), and the retroviral solution (1 mL) was added to MPI-KO HT1080 cells or MPI-KO HeLa cells (5 × 10^4^ cells/six-well plates) that were cultured for 24 h in complete medium supplemented with 20 μM Man. The transduced cells were selected for 14 days in complete medium supplemented with [for pMXs-Neo, pMXs-Neo-mCherry-hCdt1(1/100)Cy(−), and pMMLV-mVenus-hGem(1/110):IRES:Bsd] or without (pMXs-Neo-hMPI) 20 μM Man, and 600 μg/mL G418 [09380-44, Nacalai Tesque; for pMXs-Neo, pMXs-Neo-hMPI, and pMXs-Neo-mCherry-hCdt1(1/100)Cy(−)] or 13 μg/mL blasticidin S [A1113903, Thermo Fisher Scientific; for pMMLV-mVenus-hGem(1/110):IRES:Bsd].

### Preparation of whole-cell lysates

Cells were rinsed once with PBS, scraped in the same buffer, and pelleted by centrifugation at 200 × *g* for 5 min at 4°C. The wet cell weight was measured after the supernatant had been completely removed. The cell pellet was resuspended at a concentration of 10 mg wet cells/100 μL of PBS. The cell suspension was mixed with an equal volume of 2 × Laemmli sample buffer containing 10% ϕ3-mercaptoethanol, 1 × cOmplete Protease Inhibitor Cocktail (11836170001, Sigma-Aldrich), and 1 × PhosSTOP (4906845001, Sigma-Aldrich), and homogenized by two cycles of 10-s sonication at 0°C using a probe-type sonicator (VP-5S, TAITEC). The cell homogenates were denatured by heating at 100°C for 3 min and stored at −80°C until use.

### Chromatin extraction

Cells were washed once with PBS, scraped in the same buffer, and pelleted by centrifugation at 200 × *g* for 5 min at 4°C. The wet cell weight was measured after the supernatant had been completely removed. The cell pellet was extracted by incubating for 5 min at 0°C at a concentration of 10 mg wet cells/100 μL of CSK buffer [10 mM Hepes-KOH, pH7.4, 340 mM sucrose, 10% (v/v) glycerol, 10 mM KCl, 1.5 mM MgCl_2_, 0.1% (v/v) Triton X-100, 1 mM ATP, 1 mM DTT, 1 × cOmplete Protease Inhibitor Cocktail, and 1 × PhosSTOP]. The homogenate was centrifuged at 1,390 × *g* for 5 min at 4°C and the pellet was resuspended in 100 μL of CSK buffer. The homogenate was centrifuged again at 1,390 × *g* for 5 min at 4°C. The pellet was resuspended in 100 μL of CSK buffer and mixed with 100 μL of 2 × Laemmli sample buffer containing 10% β-mercaptoethanol. The samples were homogenized by two cycles of 10-s sonication at 0°C using a probe-type sonicator (VP-5S, TAITEC) and the homogenates were denatured by heating at 100°C for 3 min.

### Western and lectin blotting

Whole-cell lysates or chromatin fractions (250 μg of wet cells) were separated by sodium dodecyl sulfate (SDS)-polyacrylamide gel electrophoresis (PAGE) and analyzed by lectin blotting using biotinylated concanavalin A (Con A) lectin (1:10,000, Cosmo Bio) and VECTASTAIN ABC-HRP kit (PK-4000, Vector Laboratories) or by Western blotting using primary antibodies and secondary antibodies. The primary antibodies were as follows: anti-MPI (1:5,000, GTX103682, GeneTex), anti-β-actin (1:10,000, 010-27841, FUJIFILM Wako), anti-H2AX (1:2,000, 938CT5.1.1, Santa Cruz), anti-phospho-H2AX (Ser139) (γH2AX; 1:5,000, JBW301, Millipore), anti-RPA2 (1:5,000, 9H8, Santa Cruz), anti-Phospho-RPA2 (S33) (1:10,000, A300-246A, Bethyl), anti-MCM2 (1:10,000, D7G11, Cell Signaling Technology), anti-MCM3 (1:10,000, E-8, Santa Cruz), anti-MCM4 (1:5,000, GTX109740, GeneTex), anti-MCM5 (1:5,000, GTX33310, GeneTex), anti-MCM6 (1:20,000, H-8, Santa Cruz), anti-MCM7 (1:5,000, 141.2, Santa Cruz), anti-CDT1 (1:10,000, ab202067, Abcam), anti-CDC6 (1:1000, ab109315, Abcam), anti-ORC2 (1:10,000, 3G6, Santa Cruz), and anti-CDC45 (1:1,000, D7G6, Cell Signaling Technology). The secondary antibodies were as follows: horseradish peroxidase (HRP)-conjugated goat anti-mouse immunoglobulin (IgG) (1:10,000, 7076, Cell Signaling Technology), HRP-conjugated goat anti-rabbit IgG (1:10,000, 7074, Cell Signaling Technology), and HRP-conjugated goat anti-rat IgG (1:5,000, 7077, Cell Signaling Technology).

### Cell cycle analysis

Cells were labeled with 10 μM BrdU (B1575, Tokyo Chemical Industry) for 1 h before harvest. Trypsinized cells (1 × 10^6^ cells) were washed two times with ice-cold PBS and fixed in 1 mL of 70% ethanol for no less than 16 h at 4°C. After washing two times with PBS, fixed cells were denatured by incubating in 500 μL of 2.0 M HCl for 30 min at 25°C. Denatured cells were washed two times with PBS and once with 1 mL of sodium borate buffer, pH8.5. The cells were permeabilized in 500 μL of PBS containing 0.1% Triton X-100 (PBSTx) for 10 min at 25°C. The permeabilized cells were incubated for 1 h at 25°C in 500 μL of PBS containing 1% bovine serum albumin (BSA), Alexa Fluor 488-conjugated anti-BrdU antibody (1:100, 3D4, Biolegend), and Cellstain Hoechst 33342 solution (1:100, H342, Dojindo). The cells were then washed once with 1 mL of PBS containing 1% BSA (PBS/BSA) and resuspended in 500 μL of the same buffer. The stained samples were analyzed by BD LSRFortessa X-20 and FlowJo v10.6 (Becton Dickinson).

### Chromatin flow cytometry

Trypsinized cells (1 × 10^6^ cells) were extracted by incubating for 5 min at 0°C in 500 μL of CSK buffer. The cells were washed by adding 1 mL of ice-cold PBS/BSA, pelleted by centrifugation at 1,390 × *g* for 5 min at 4°C, and fixed for 15 min at 25°C in 500 μL of 4% paraformaldehyde (161-20141, FUJIFILM Wako). After quenching the reaction by adding 1 mL of PBS/BSA, the cells were pelleted by centrifugation at 2,000 × *g* for 7 min at 4°C and washed again with 1 mL of PBS/BSA. The washed cells were then resuspended in 200 μL of PBS containing 1% BSA and 0.1% Triton X-100 (PBSTx/BSA) containing primary antibodies [MCM2 (1:200, D7G11, Cell Signaling Technology) and γH2AX (1:200, JBW301, Millipore)] and incubated for 1 h at 25°C. After washing the cells once in 1 mL of PBSTx/BSA, they were incubated for 1 h at 25°C in the dark with 200 μL of PBSTx/BSA containing secondary antibodies [R37118, donkey anti-rabbit IgG-Alexa Fluor 488 (1:1,000, Thermo Fisher Scientific); A-31571, donkey anti-mouse IgG-Alexa Fluor 647 (1:1,000, Thermo Fisher Scientific)] and Cellstain Hoechst 33342 solution (1:200). The cells were washed once in 1 mL of PBSTx/BSA, resuspended in 500 μL of the same buffer, and subjected to flow cytometry.

### Cell viability (ATP) assay

Cells (1 × 10^3^ cells/96-well plate) were seeded and cultured for 24 h in 100 μL of complete medium supplemented with 20 μM Man. For co-treatment assay, the cells were incubated with drugs or vehicle alone for 24 h in complete medium supplemented with 50 μM or 5 mM Man. For a preconditioning assay, cells were incubated for 24 h in complete medium supplemented with 50 μM or 5 mM Man, and the preconditioned cells were further incubated with drugs or vehicle alone for 24 h. After the drug treatment, CellTiter-Glo 2.0 (100 μL/well, Promega) was added to the cells and incubated for 10 min at 25°C before measuring luminescence with an integration time of 1,000 ms using an Infinite 200 Pro microplate reader (Tecan).

### LDH assay

For Fig. 5E, cells (1 × 10^3^ cells/96-well plate) that were cultured for 24 h in 100 μL of complete medium supplemented with 20 μM Man were treated with 30 nM IACS-010759 or dimethylsulfoxide (DMSO) alone for an additional 6 h in complete medium supplemented with unchallenged or challenged concentrations of Man. Total LDH activity was measured using Cytotoxicity LDH Assay Kit-WST (Dojindo), in accordance with the manufacturer’s instructions. For Fig. 5F (necrosis assay), cells (3.3 × 10^4^ cells/24-well plate) that were cultured for 24 h in 500 μL of complete medium supplemented with 20 μM Man were treated with 30 nM IACS-010759 or DMSO alone for an additional 24 h in 500 μL of complete medium supplemented with unchallenged or challenged concentrations of Man. After centrifugation at 250 × *g* for 5 min at 4°C, the culture supernatant (50 μL, extracellular LDH) was transferred to a 96-well plate. The cells were solubilized by adding 20 μL of 10% (w/v) Tween 20 for 30 min at 37°C with gentle agitation. The homogenates (50 μL, total LDH) were transferred to the same 96-well plate prepared as above. LDH activity was measured using Cytotoxicity LDH Assay Kit-WST, in accordance with the manufacturer’s instructions.

### Seahorse real-time cell metabolic analysis

The ATP rate assay was performed using an XFe96 Extracellular Flux analyzer, in accordance with the manufacturer’s instructions (Seahorse Bioscience). MPI-KO HT1080 cells (#3, 1 × 10^4^ cells/96-well assay plate) were seeded and incubated for 18 h in complete DMEM. Prior to the assay, the cells were preincubated for 60 min at 37°C in XF DMEM medium containing 10% FBS, 10 mM glucose, 4 mM L-glutamine, and 50 μM Man. The cells were then treated with 5 mM Man or medium alone for 120 min, followed by incubation with 1.5 μM oligomycin for 18 min and then 0.5 μM each of rotenone and antimycin A for 18 min. Data analysis was performed using the XF Real-Time ATP Rate Assay Report Generator (Seahorse Bioscience).

### Label-free proteomics

Cells (5 × 10^5^ cells/10-cm dish) were cultured for 24 h in 10 mL of complete medium supplemented with 20 μM Man. The cells were further incubated for 1, 2, and 6 days in 10 mL of complete medium supplemented with 50 μM Man (1 and 2 days) or 5 mM Man (1, 2, and 6 days). For 6-day incubation, the medium was replaced every other day. The cells were washed two times with PBS and scraped in 1 mL of PBS. After the cells had been pelleted by centrifugation at 200 × *g* for 5 min at 4°C, the cell pellets were flash-frozen in liquid nitrogen and stored at −80°C until use. Protein precipitates were obtained by adding 10% trichloroacetic acid to freeze-thawed cells and centrifuging at 12,000 × *g* for 20 min. After washing the precipitates three times with acetone, they were dissolved with 7 M guanidine, 1 M Tris-HCl (pH 8.5), 10 mM EDTA, and 50 mM dithiothreitol. After alkylation with iodoacetic acid, samples were desalted using PAGE Clean Up Kit (Nacalai Tesque). The resultant precipitates were dissolved with 20 mM Tris-HCl (pH 8.0), 0.03% (w/v) n-dodecyl-β-D-maltoside, and digested with trypsin (tosyl phenylalanyl chloromethyl ketone-treated; Worthington Biochemical) at 37°C for 18 h. The concentration of the peptide mixture was quantified by amino acid analysis (Masuda and Dohmae, 2011). One microgram of each peptide mixture was subjected to liquid chromatography (LC)-mass spectrometry (MS)/MS. Solvent A (0.1% formic acid) and solvent B (80% acetonitrile with 0.1% formic acid) were used as eluents. Peptides were separated using an Easy nLC 1200 (Thermo Fisher Scientific) equipped with a nano-ESI spray column (NTCC-360, 0.075 mm internal diameter × 105 mm length, 3 μm, Nikkyo Technos) at a flow rate of 300 nl/min under linear gradient conditions over 250 min. The separated peptides were analyzed with an online coupled Q Exactive HF-X Mass Spectrometer (Thermo Fisher Scientific) using the data-dependent Top 10 method. The acquired data were processed using MASCOT 2.8 (Matrix Science) and Proteome Discoverer 2.4 (Thermo Fisher Scientific). The MASCOT search was conducted as follows: Database, NCBIprot; taxonomy, *Homo sapiens* (human) (438,061 sequences); type of search, MS/MS ion; enzyme, trypsin; fixed modification, none; variable modifications, acetyl (protein N-term), Gln->pyro-Glu (N-term Q), oxidation (M), carboxymethyl (C); mass values, monoisotopic; peptide mass tolerance, ± 15 ppm; fragment mass tolerance, ± 30 mmu; max. missed cleavages, 3; and instrument type, ESI-TRAP. Label-free quantification was performed using the quantification method based on the ion intensity of peptides in Proteome Discoverer 2.4.

### Metabolomics

Approximately 8 × 10^6^ cells were washed two times with 5 mL of 5% (w/v) mannitol (133-00845, FUJIFILM Wako) and scraped in 1.3 mL of methanol (138-14521, FUJIFILM Wako) that was spiked with 10 μM external standards (Human Metabolome Technologies). The cell homogenates were spun at 15,000 × *g* for 5 min at 4°C and the wet cell weights were measured. The supernatant was analyzed by capillary electrophoresis (CE) time-of-flight (TOF) mass spectrometry (MS) using an Agilent CE-TOFMS system (Agilent Technologies) at Human Metabolome Technologies.

### Time-lapse imaging

Fucci MPI-KO HT1080 cells (#3 subclone 9-5, 4 × 10^4^ cells/six-well dish) were cultured for 24 h in 2.5 mL of complete medium supplemented with 20 μM Man. The cells were washed once with complete DMEM with no phenol red (040-30095, FUJIFILM Wako), 4 mM glutamine, and 10% FBS. The washed cells were cultured in 2.5 mL of the same medium supplemented with 50 μM Man or 5 mM Man in a humidified chamber (Tokai Hit) at 37°C with 5% CO_2_. Fluorescence and differential interference contrast images were obtained every 15 min using KEYENCE BZ-X810 with a PlanFluor 20× objective lens (NA=0.45, WD=8.80-7.50 mm, Ph1; KEYENCE). Fucci mCherry and mVenus signals were detected using a TRITC filter (Ex, 545/25 nm; Em, 605/70 nm; KEYENCE) and mVenus filter (Ex, 500/20; Em, 535/30 nm; M SQUARE), respectively. Images were acquired after a 30-min equilibration period.

### Quantification

For Western blots, the X-ray films were scanned using an EPSON GT-7600UF scanner. ImageJ (Schneider et al., 2012) was used for the quantification of band intensity. For image processing of Fucci time-lapse imaging, a Fiji/ImageJ plugin, Trackmate (Tinevez et al., 2017), was used to track single cells. MATLAB (MathWorks, Natick, MA) was used to quantify the duration of each cell cycle phase. The Fucci profiles in the mannose-challenged cells (Fig. 2E, fig. S4) were visually inspected for classification.

### Statistical analysis

R version 3.3.3 and Prism 9 were used for statistical analysis. Statistical analysis was performed by applying an unpaired two-sided Welch’s *t*-test for comparison of the means between two groups. Comparisons of the means among more than two groups were performed with one-way or two-way analysis of variance (ANOVA) followed by *post hoc* testing with Dunnett’s test or Tukey’s test. Data are reported as the mean ± standard deviation (SD). *P*-values < 0.05 were considered to be statistically significant.

## Acknowledgments

We wish to thank our laboratory members, Drs. Takashi Akazawa and Yosuke Matsuoka (Osaka International Cancer Institute), for fruitful discussions, Dr. Steven. A. Rosenberg for MCA205 cells, and Drs. Kazuki Nakajima (Gifu University) and Takuro Tojima (RIKEN) for technical advice. We thank Edanz (https://jp.edanz.com/ac) for editing a draft of this manuscript.

## Funding

The Takeda Science Foundation (to YH)

The Rocket Fund (to HHF)

R01DK99551 (to HHF)

## Author contributions

Conceptualization: YH

Methodology: YH, TH, HHF

Investigation: YH, YM, TS, MN, AU, YI, KM, YO

Funding acquisition: YH

Project administration: YH

Supervision: JM, EM, SH, HU, HT, NT

Writing – original draft: YH, YM, TS, MN, YI

Writing – review & editing: All authors reviewed and approved final manuscript.

## Data and materials availability

The mass spectrometry proteomics data have been deposited to the ProteomeXchange Consortium via the PRIDE partner repository with the dataset identifier PXD036449 and 10.6019/PXD036449 (Username; Password: aRfYzEvX).

## Supplementary Materials

figs. S1 to S8

tables S1 to S2

References (1–45)

movies S1 to S2

data S1 to S3

## Supplementary Materials

### This PDF file includes

figs. S1 to S8

Tables S1 to S2

Captions for movies S1 to S2

Captions for Data S1 to S3

### Other Supplementary Materials for this manuscript include the following

movies S1 to S2

Data S1 to S3 [S1. Proteomic data], [S2. Functional annotation analysis of proteomic data], [S3. Metabolomic data]

**fig. S1.**
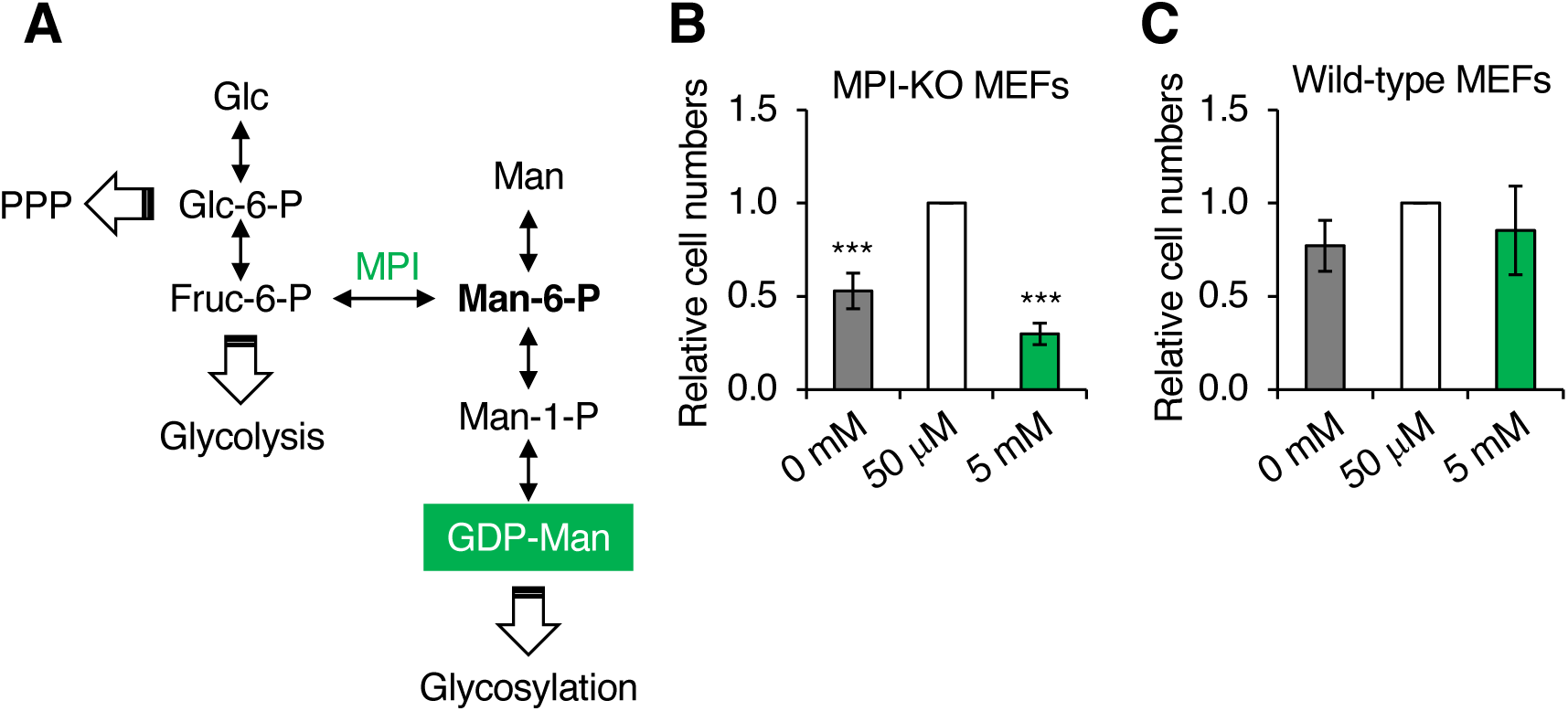
Mannose auxotrophy and sensitivity in MPI-KO MEFs and wild-type MEFs. (**A**) Metabolic pathways for glucose and mannose. PPP, pentose phosphate pathway; Glc, glucose; Glc-6-P, glucose-6-phosphate; Fruc-6-P, fructose-6-phosphate; Man-6-P, mannose-6-phosphate; Man-1-P, mannose-1-phosphate; Man, mannose; GDP-Man, guanosine diphosphate-mannose. (**B** and **C**) The relative numbers of MPI-KO MEFs (**B**) and wild-type MEFs (**C**) after 48-h incubation in culture medium supplemented with mannose at the indicated concentrations. Data represent the mean ± SD; n = 3 independent experiments. \*\*\**p* < 0.001, one-way ANOVA with post hoc Dunnett’s test (**B** and **C**).

**fig. S2.**
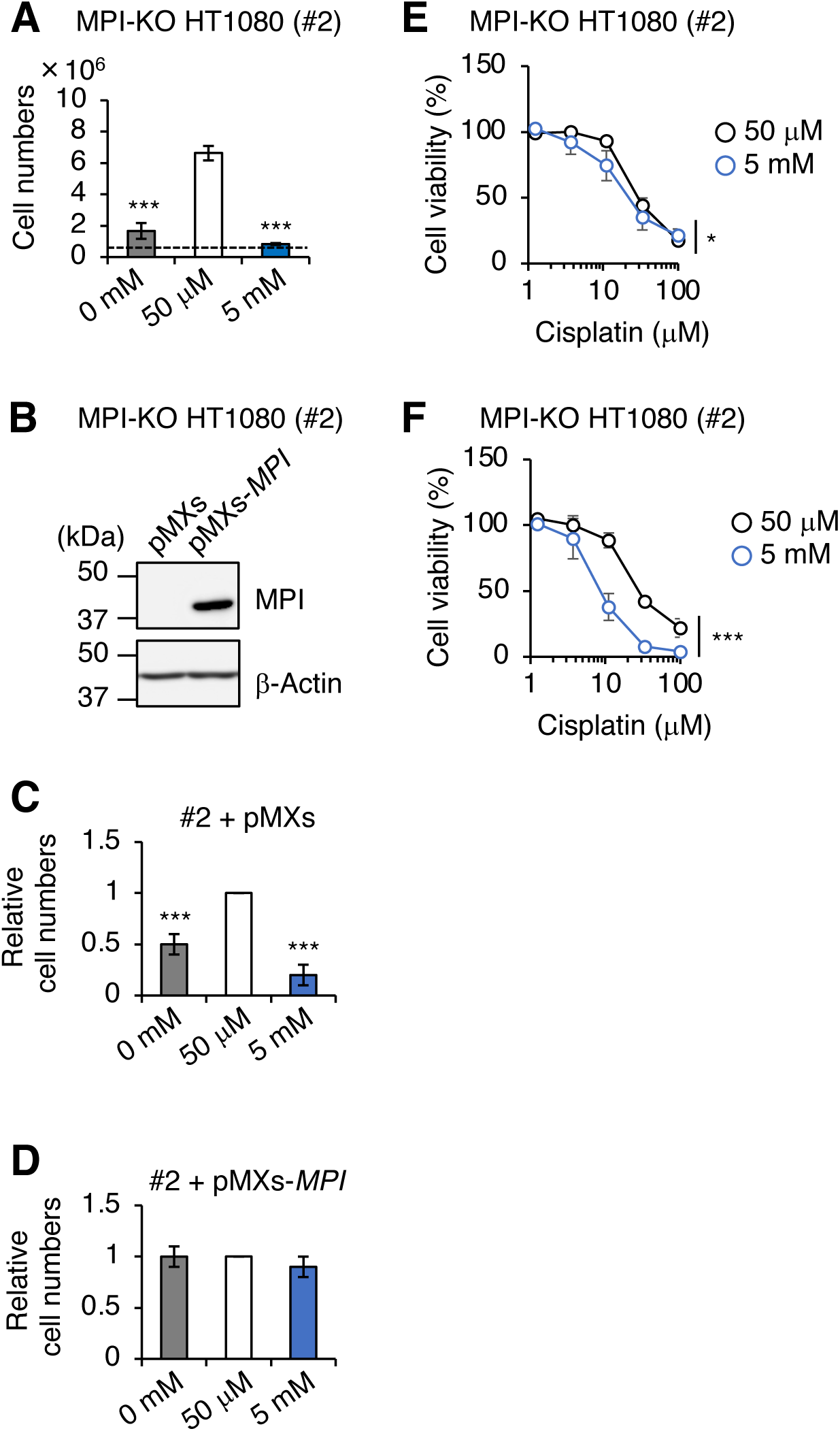
Establishment of MPI-KO HT1080 cells. (**A**) The numbers of MPI-KO HT1080 (#2) cells after 48-h incubation in culture medium under the indicated conditions. A dashed line indicates the cell numbers at seeding. (**B**) Western blot analysis of MPI-KO HT1080 (#2) cells retrovirally transduced with empty vector (pMXs) or human MPI gene (pMXs-*MPI*). (**C** and **D**) The relative numbers of MPI-KO HT1080 (#2) cells retrovirally transduced with empty vector (pMXs, **C**) or human *MPI* gene (pMXs-*MPI*, **D**) after 48-h incubation in culture medium supplemented with mannose at the indicated concentrations. (**E** and **F**) Cell viability assay in MPI-KO HT1080 (#2) cells co-treated with mannose (50 μM or 5 mM) and cisplatin for 24 h (**E**), or preconditioned with mannose (50 μM or 5 mM) for 24 h, followed by incubation with cisplatin for an additional 24 h (**F**). Data represent the mean ± SD; n = 3 independent experiments. \**p* < 0.05 and \*\*\**p* < 0.001, one-way ANOVA with post hoc Dunnett’s test (**A**, **C**, and **D**) or two-way ANOVA (**E** and **F**).

**fig. S3.**
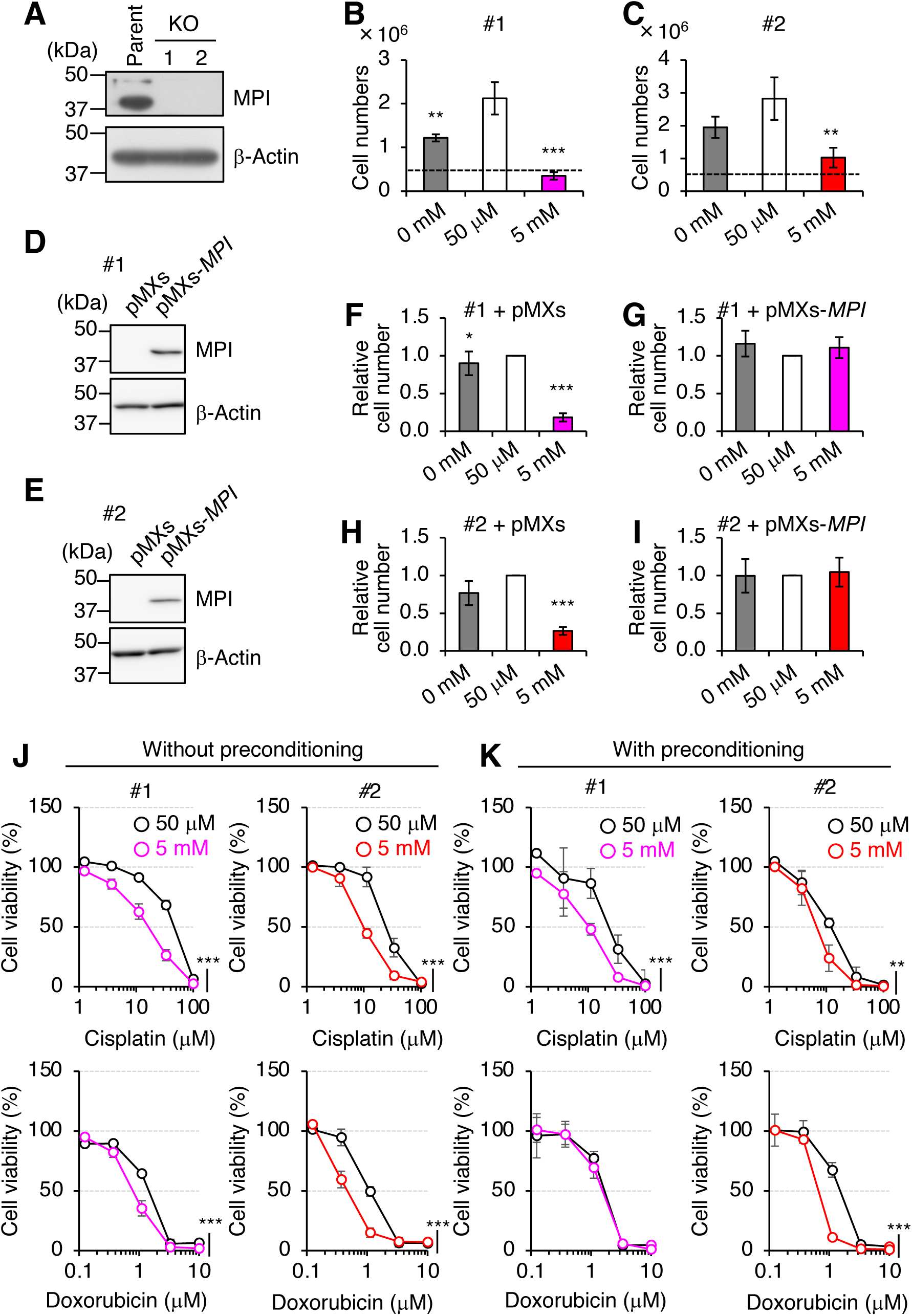
Establishment of MPI-KO HeLa cells. (**A**) Western blot analysis of HeLa (parent) cells and MPI-KO HeLa (KO, clones #1 and #2) cells. (**B** and **C**) The numbers of MPI-KO HeLa (#1, **B**; and #2, **C**) cells after 48-h incubation under the indicated conditions. (**D** and **E**) Western blot analysis of MPI-KO HeLa (#1, **D**; and #2, **E**) cells retrovirally transduced with empty vector (pMXs) or human *MPI* gene (pMXs-*MPI*). (**F**–**I**) The relative numbers of MPI-KO HeLa (#1, **F** and **G**; and #2, **H** and **I**) cells retrovirally transduced with empty vector (pMXs, **F** and **H**) or human *MPI* gene (pMXs-*MPI*, **G** and **I**) after 48-h incubation under the indicated conditions. (**J** and **K**) Cell viability assay in MPI-KO HeLa (#1 and #2) cells co-treated with mannose (50 μM or 5 mM) and DNA replication inhibitors (cisplatin or doxorubicin) for 24 h (without preconditioning, **J**), or preconditioned with mannose (50 μM or 5 mM) for 24 h, followed by incubation with the DNA replication inhibitors for an additional 24 h (with preconditioning, **K**). Data represent the mean ± SD; n = 3 independent experiments. \**p* < 0.05, **p < 0.01, and ***p < 0.001, one-way ANOVA with post hoc Dunnett’s test (**B**, **C**, and **F**–**I**) or two-way ANOVA (**J** and **K**).

**fig. S4.**
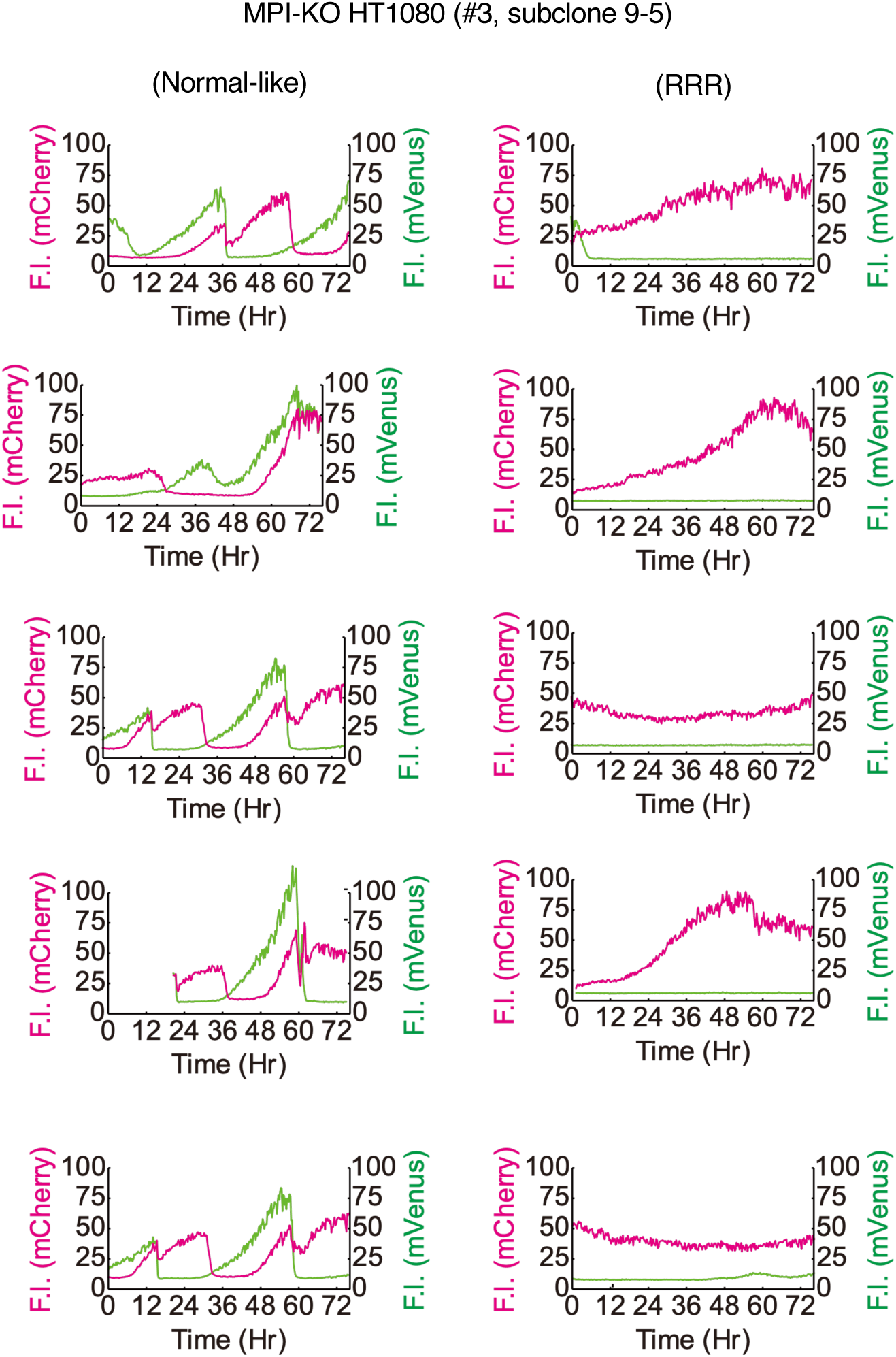
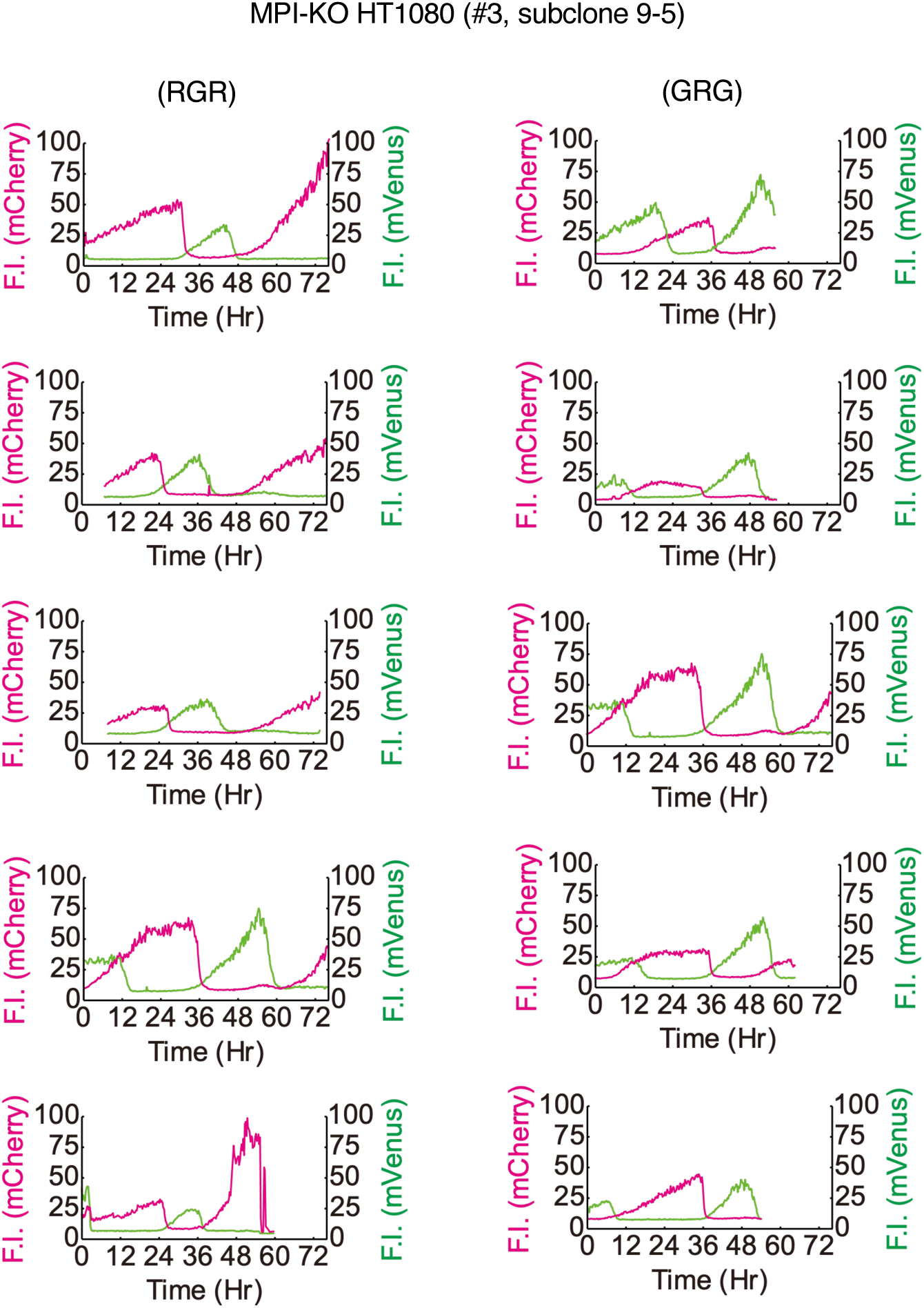
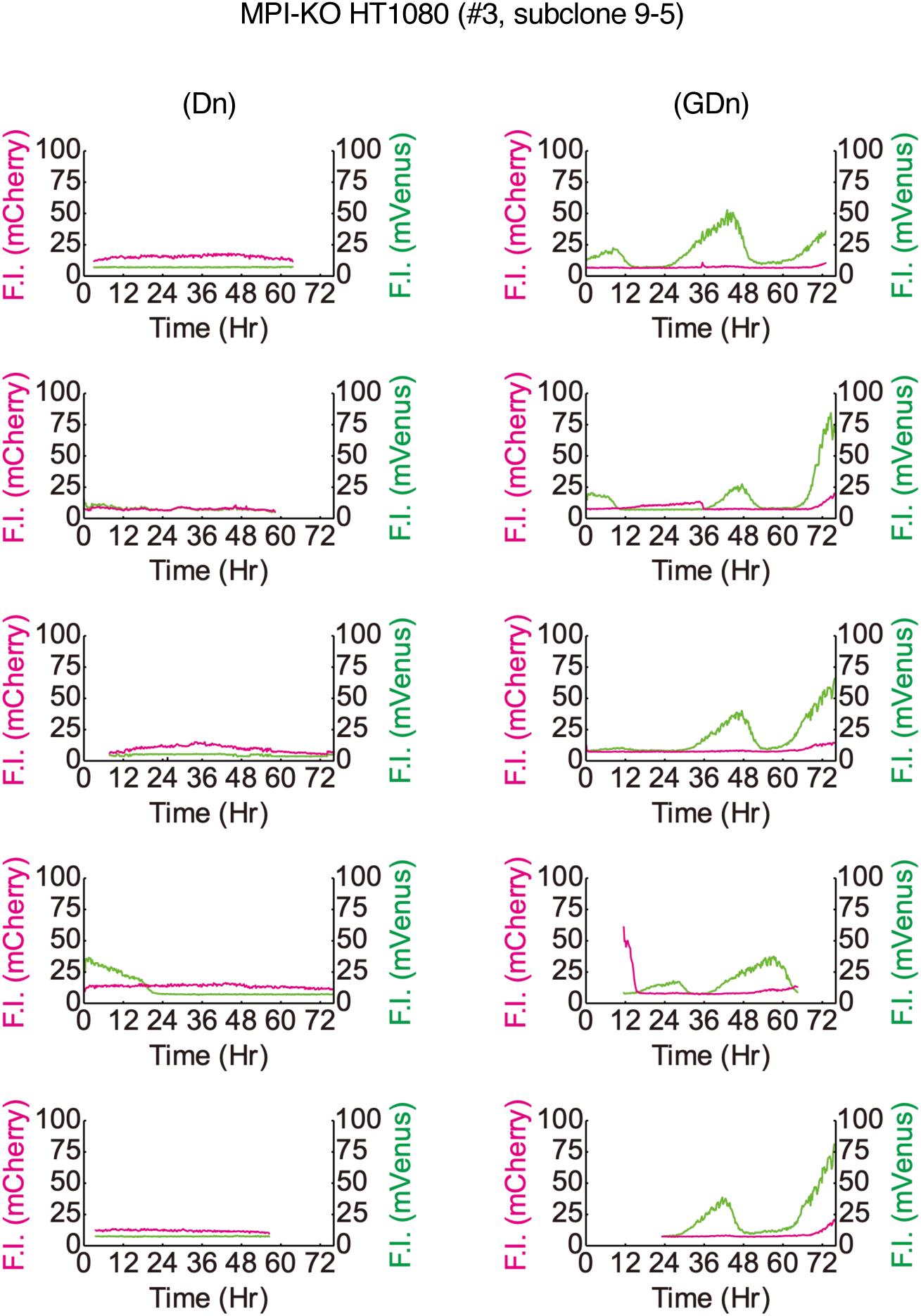
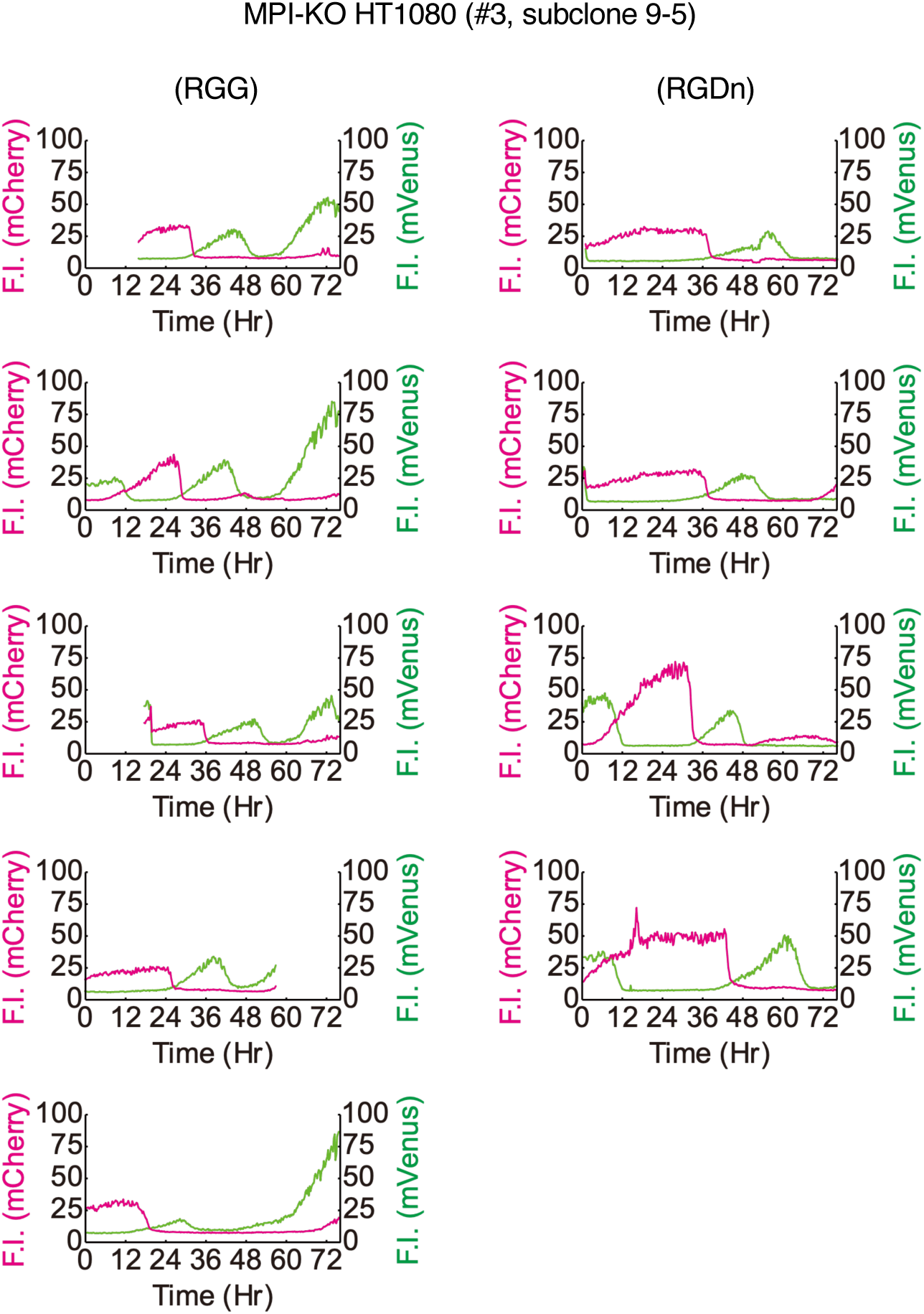
The classification of Fucci profiles in MPI-KO HT1080 cells cultured under mannose-challenged conditions. Fluorescent intensity (F.I.) for mCherry (left y-axis) and mVenus (right y-axis) is shown in magenta and green, respectively. The order of expression of mCherry-hCdt1(1/100)Cy(–) (denoted by R) and mVenus-hGem(1/110) (denoted by G) was used to classify the Fucci profiles and the classification is indicated in parentheses. Dn, double negative for Fucci indicators; Not clear, Fucci profiles that could not be classified into categories defined in this study. One hundred cells were visually inspected for the classification and five examples are shown for each category, except for RGDn (n = 4 in total). Cells that could not be classified visually were categorized as “not clear.”

**fig. S5.**
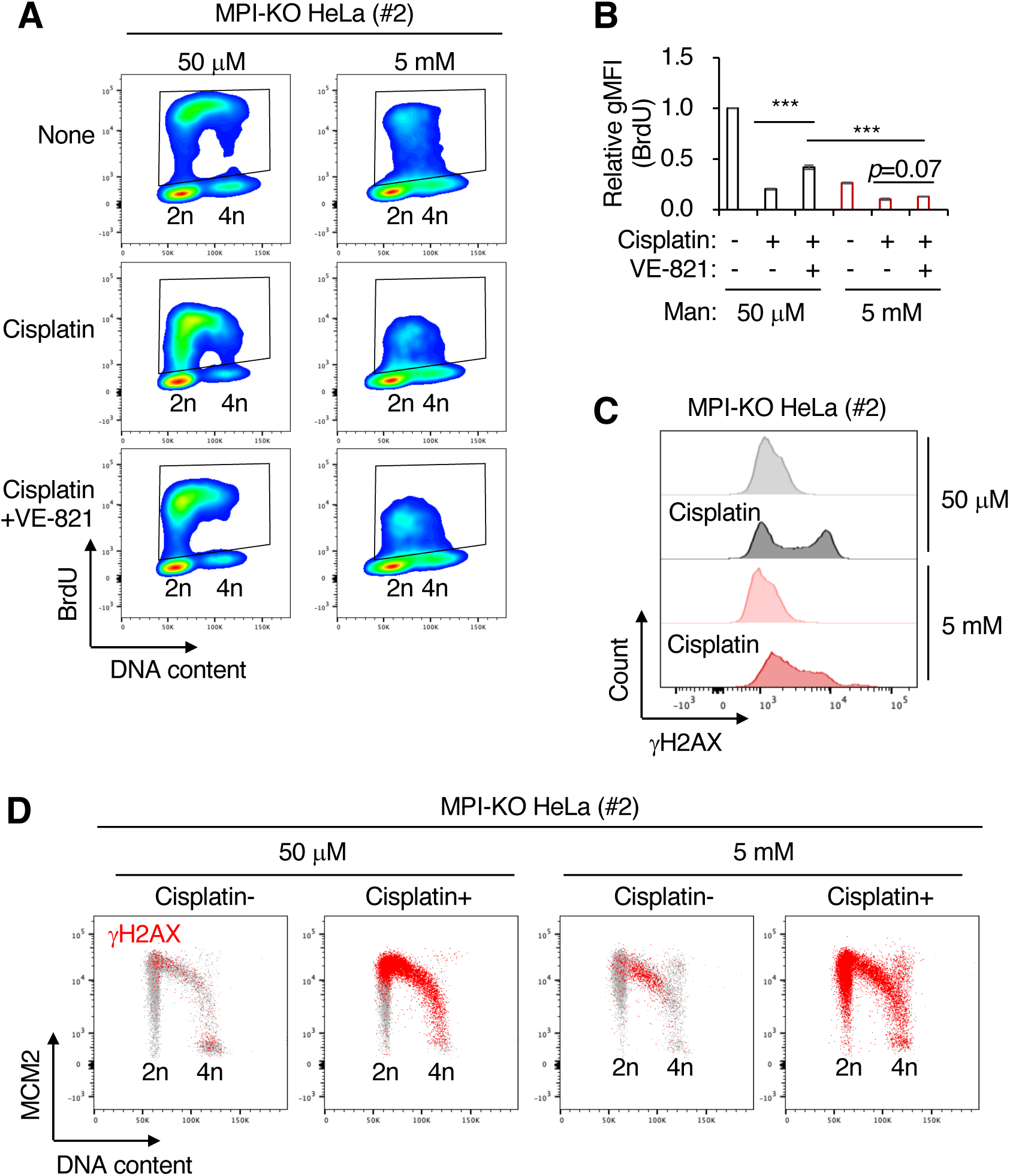
Mannose challenge disengages dormant origins from DNA synthesis during replication stress in MPI-KO HeLa cells. (**A**) Flow cytometry for BrdU and DNA content in MPI-KO HeLa (#2) cells treated as in Fig. 4A. (**B**) Relative geometric mean fluorescent intensity (gMFI) of BrdU in **B** (in the gated populations). (**C** and **D**) Chromatin flow cytometry for γH2AX, MCM2, and DNA content (Hoechst33342) in MPI-KO HeLa (#2) cells treated as in **A**, except that VE-821 treatment and BrdU labeling were omitted. Data represent the mean ± SD; n = 3 independent experiments. \*\*\**p* < 0.001, one-way ANOVA with post hoc Tukey’s test (**B**).

**fig. S6.**
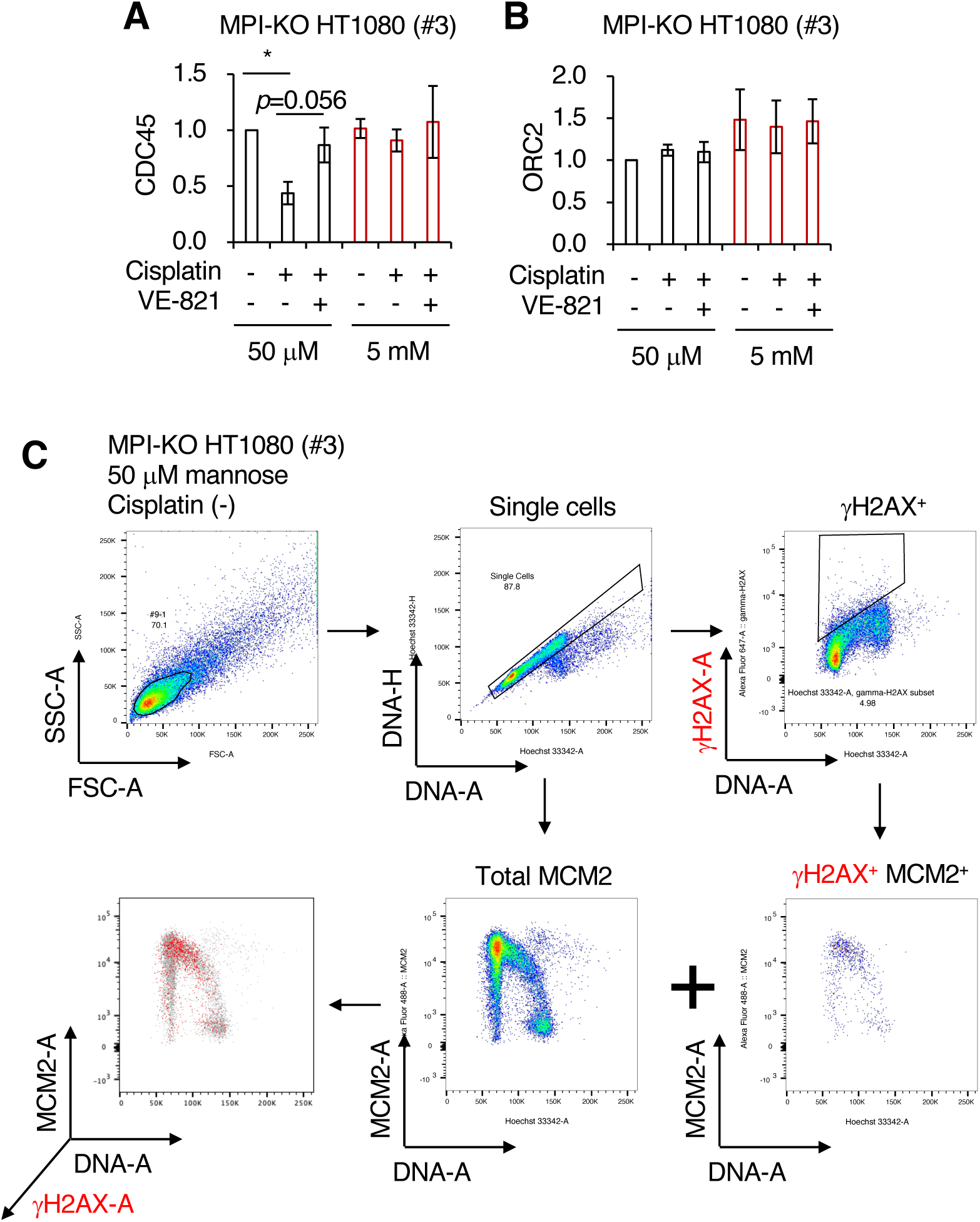
Mannose challenge disengages dormant origins from DNA synthesis during replication stress in MPI-KO HT1080 cells. (**A and B**) Relative amounts of the chromatin-bound CDC45 (A) and ORC2 (B) in Fig. 4D. (**C**) Strategy for gating the chromatin-bound γH2AX and MCM2 in Fig. 4F. Data represent the mean ± SD; n = 3 independent experiments. \**p* < 0.05, one-way ANOVA with post hoc Tukey’s test (A and B).

**fig. S7.**
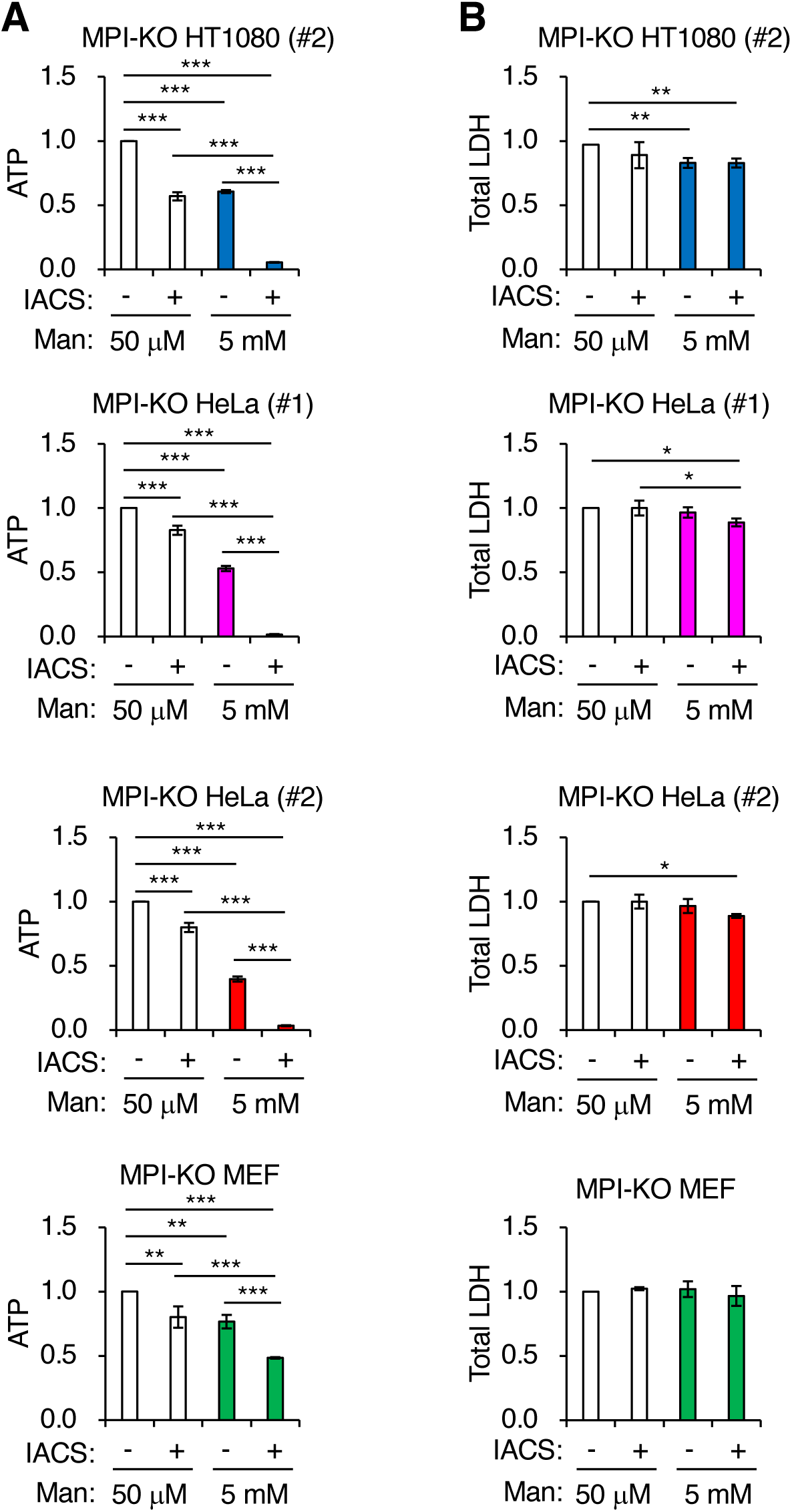
Mannose-challenged MPI-KO cells highly depend on OXPHOS for ATP production. Relative ATP levels (**A**) and LDH activity (**B**) in MPI-KO HT1080 (#2) cells, MPI-KO HeLa (#1 and #2) cells, and MPI-KO MEFs co-treated with mannose (50 μM or 5 mM) and 30 nM IACS-010759 (IACS) or vehicle alone for 6 h. Data represent the mean ± SD; n = 3 independent experiments. \**p* < 0.05, \*\**p* < 0.01, and \*\*\**p* < 0.001, one-way ANOVA with post hoc Tukey’s test.

**fig. S8.**
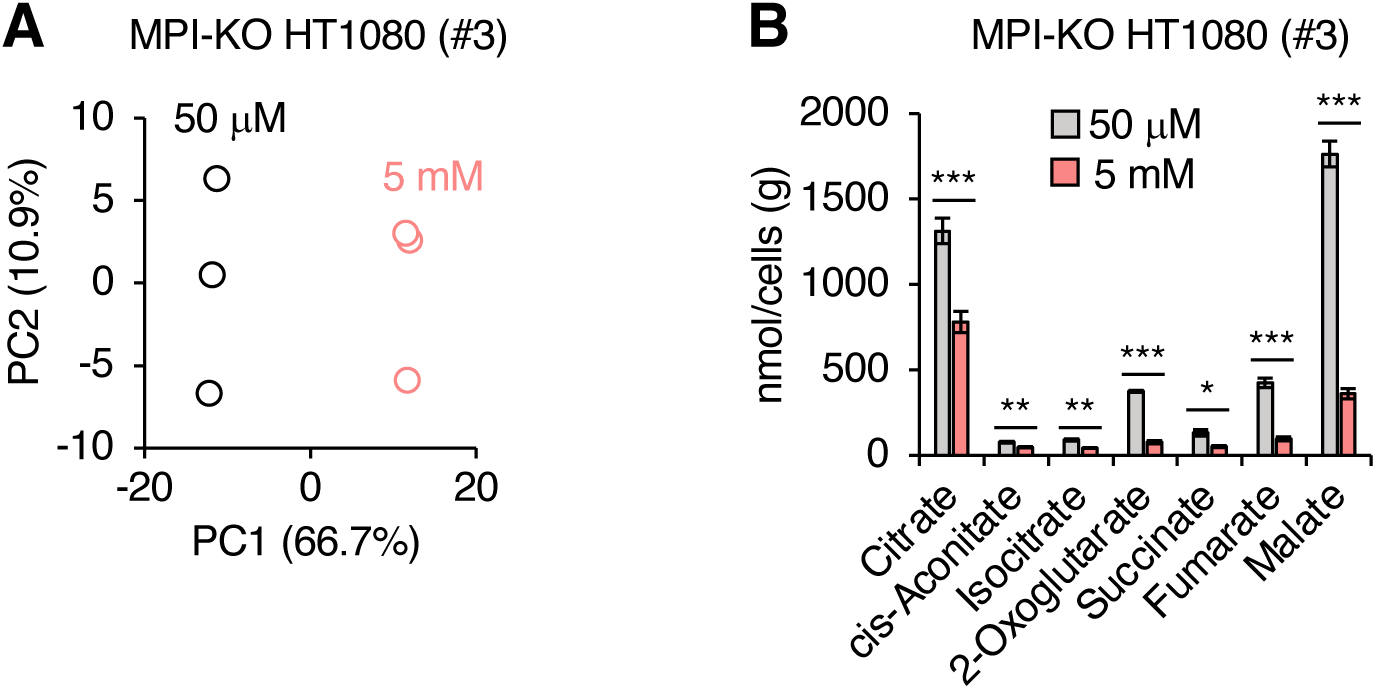
Metabolomics of mannose-challenged and -unchallenged MPI-KO HT1080 cells. (**A**) Principal component analysis of metabolites in MPI-KO HT1080 (#3) cells cultured in the presence of 50 μM or 5 mM mannose for 24 h. (**B**) Quantification of metabolites in tricarboxylic acid cycle. Data represent the mean ± SD; n = 3 independent experiments. \**p* < 0.05, \*\**p* < 0.01, \*\*\**p* < 0.001, Welch’s *t*-test.

**Table S1.**
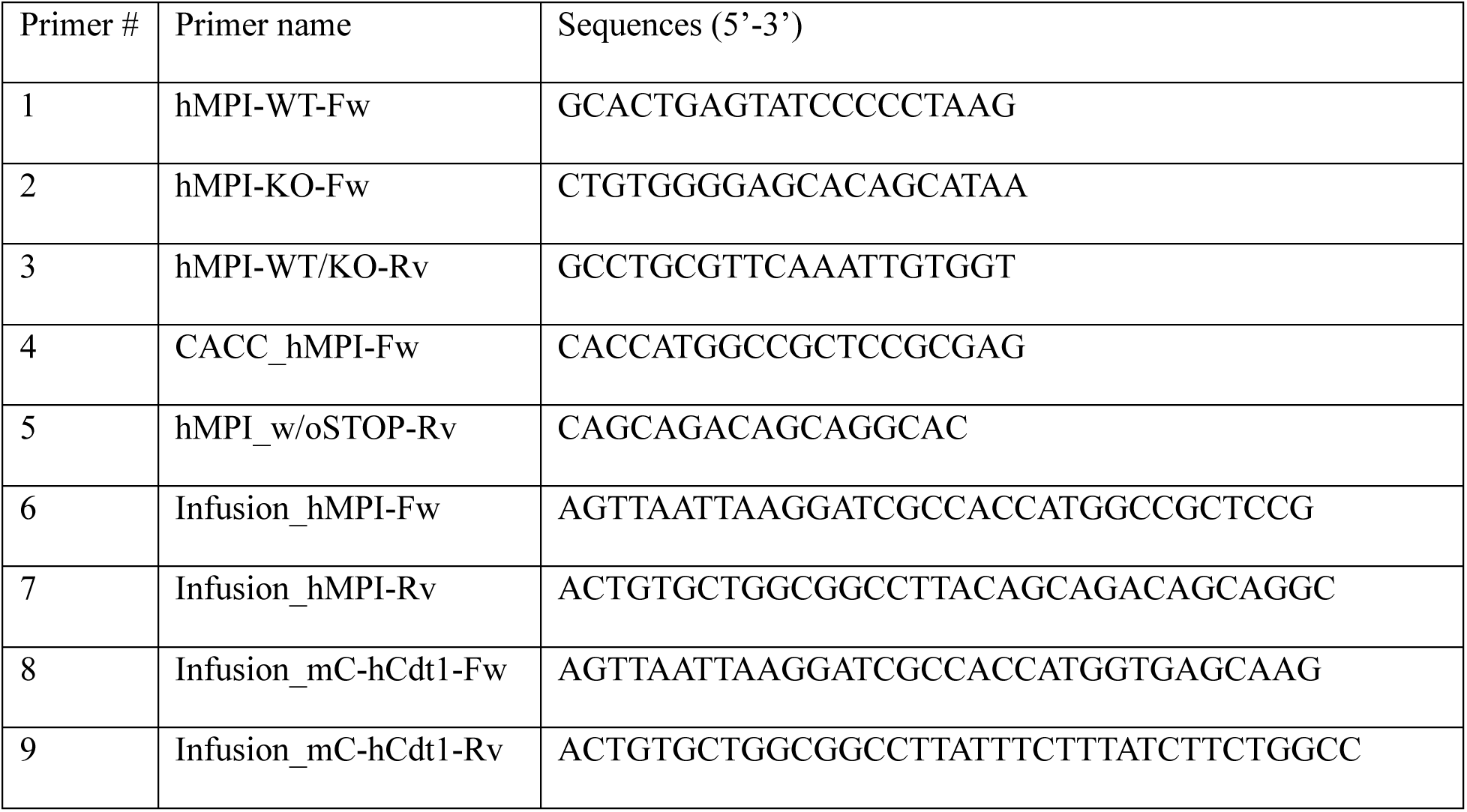
List of primers used in this study.

**Table S2.** List of plasmids used in this study.

| Plasmid # | Plasmid name | Notes |
| --- | --- | --- |
| 1 | pENTR-hMPI | This study |
| 2 | pMXs-Neo-hMPI | This study |
| 3 | mCherry-hCdt1(1/100)Cy(-)/pcDNA3 | (Sakaue-Sawano *et al.*, 2017) |
| 4 | pMXs-Neo-mCherry-hCdt1(1/100)Cy(-) | This study |
| 5 | pMMLV-mVenus-hGem(1/110):IRES:Bsd | This study |

**movie S1. Time-lapse imaging of cell cycles in MPI-KO HT1080.**

MPI-KO HT1080 that expressed Fucci(CA) reporters was observed for 76 hours under the mannose-unchallenged (left) or the mannose-challenged (right) culture conditions. Red, mCherry-hCdt1(1/100)Cy(-); Green, mVenus-hGem(1/110). Images were acquired every 15 min. Image size: 253.44 μm × 253.44 μm for each panel.

**movie S2. Abnormal cytokinesis during M phase in the mannose-challenged MPI-KO HT1080 with normal-like Fucci(CA) signal profiles.**

Time-lapse movies of three representative Fucci(CA)-expressing MPI-KO HT1080 that were cultured under the mannose challenge conditions. Red, mCherry-hCdt1(1/100)Cy(-); Green, mVenus-hGem(1/110). Note that the cells in G2-M phase (yellow) show abnormal cytokinesis. Images were acquired every 15 min. Image size: 52.8 μm × 52.8 μm for each panel.

**Data S1. (separate file)**

Proteomic data.

**Data S2. (separate file)**

Functional annotation analysis of proteomic data.

**Data S3. (separate file)**

Metabolomic data.

